# Establishing the Validity of Compressed Sensing Diffusion Spectrum Imaging

**DOI:** 10.1101/2023.02.22.529546

**Authors:** Hamsanandini Radhakrishnan, Chenying Zhao, Valerie J. Sydnor, Erica B. Baller, Philip A. Cook, Damien Fair, Barry Giesbrecht, Bart Larsen, Kristin Murtha, David R. Roalf, Sage Rush-Goebel, Russell Shinohara, Haochang Shou, M. Dylan Tisdall, Jean Vettel, Scott Grafton, Matthew Cieslak, Theodore Satterthwaite

**Affiliations:** Lifespan Informatics and Neuroimaging Center, University of Pennsylvania, Philadelphia, PA, USA; Department of Psychiatry, Perelman School of Medicine, University of Pennsylvania, Philadelphia, PA, USA; Lifespan Brain Institute, Perelman School of Medicine, University of Pennsylvania, Philadelphia, PA, USA; Department of Bioengineering, School of Engineering and Applied Science, University of Pennsylvania, Philadelphia, PA, USA; Department of Radiology, Perelman School of Medicine, University of Pennsylvania, Philadelphia, PA, USA; Masonic Institute for the Developing Brain, University of Minnesota, Minneapolis, MN, USA; Department of Psychological and Brain Sciences, University of California, Santa Barbara, CA, USA; Department of Biostatistics, Epidemiology and Informatics, University of Pennsylvania, Philadelphia, PA, USA; Center for Biomedical Image Computing & Analytics, University of Pennsylvania, Philadelphia, PA, USA; U.S. Army Research Laboratory, Aberdeen Proving Ground, MD, USA

**Author notes:** Please direct correspondence to: Theodore Satterthwaite, Richards Medical Labs, 5th Floor, Pod A, 3700 Hamilton Walk, Philadelphia, PA 19104. denotes equal contribution (Matthew Cieslak, Theodore Satterthwaite).

**Keywords:** MRI acquisition, compressed sensing, diffusion-weighted imaging, white matter

## Abstract

Diffusion Spectrum Imaging (DSI) using dense Cartesian sampling of *q*-space has been shown to provide important advantages for modeling complex white matter architecture. However, its adoption has been limited by the lengthy acquisition time required. Sparser sampling of *q*-space combined with compressed sensing (CS) reconstruction techniques has been proposed as a way to reduce the scan time of DSI acquisitions. However prior studies have mainly evaluated CS-DSI in post-mortem or non-human data. At present, the capacity for CS-DSI to provide accurate and reliable measures of white matter anatomy and microstructure in the living human brain remains unclear. We evaluated the accuracy and inter-scan reliability of 6 different CS-DSI schemes that provided up to 80% reductions in scan time compared to a full DSI scheme. We capitalized on a dataset of twenty-six participants who were scanned over eight independent sessions using a full DSI scheme. From this full DSI scheme, we subsampled images to create a range of CS-DSI images. This allowed us to compare the accuracy and inter-scan reliability of derived measures of white matter structure (bundle segmentation, voxel-wise scalar maps) produced by the CS-DSI and the full DSI schemes. We found that CS-DSI estimates of both bundle segmentations and voxel-wise scalars were nearly as accurate and reliable as those generated by the full DSI scheme. Moreover, we found that the accuracy and reliability of CS-DSI was higher in white matter bundles that were more reliably segmented by the full DSI scheme. As a final step, we replicated the accuracy of CS-DSI in a prospectively acquired dataset (n=20, scanned once). Together, these results illustrate the utility of CS-DSI for reliably delineating *in vivo* white matter architecture in a fraction of the scan time, underscoring its promise for both clinical and research applications.

## 1 Introduction

Diffusion-weighted Magnetic Resonance Imaging (dMRI) has become the dominant imaging modality for non-invasively characterizing white matter architecture in humans, with myriad applications in both research and clinical practice. While a large and growing variety of dMRI sequences exist, they all sample three spatial dimensions in *k*-space and three additional dimensions in *q*-space, which is defined by the diffusion-encoding gradients (Callaghan et al., 1988). Each 3D volume in a dMRI sequence samples a coordinate in *q*-space, with the entire set of points defining a *q*-space sampling scheme. The dMRI field has explored a wide range of *q*-space sampling schemes (Menzel et al., 2011a; V.J. Wedeen et al., 2005), with most sampling points on the surface of one or more spheres (aka “shells”). One alternative strategy densely samples *q*-space on a Cartesian grid, enabling the direct estimation of the diffusion Ensemble Average Propagator (EAP), the physical process driving biologically meaningful derivatives of dMRI. This scheme is often called Diffusion Spectrum Imaging (DSI) and can resolve crossing fibers and other complex tract architectures. However, Cartesian sampling requires long scan times, which has limited the use of DSI in translational research or clinical practice. Recently, sparse random sampling of *q*-space combined with compressed sensing (CS) reconstruction techniques, has shown massive promise in both post-mortem and animal studies (Jones et al., 2021; Menzel et al., 2011b; Naeyaert et al., 2021; Paquette et al., 2015). However, the validity of common white matter derivatives generated by compressed sensing accelerated DSI (CS-DSI) has not been adequately tested in *in vivo* human data.. Here, we sought to validate the ability of CS-DSI to characterize both macro- and micro-scale white matter properties in living humans by evaluating its accuracy and reliability compared to a full DSI scheme (which sampled 258 points in *q*-space).

The Cartesian sampling schemes used in DSI provide several important practical and theoretical advantages due to their ability to model the EAP directly. DSI allows direct measurement of the EAP in each voxel due to the Fourier relationship between *q*-space and the EAP (Van J. Wedeen et al., 2005). This relationship between the *q*-space signal and the EAP obviates the choice of which shells to sample (i.e., which *b*-values for each shell).For example, high *b*-value shells can be more useful for tractography in the presence of crossing fibers while low *b-*value shells are required for estimating some free water parameters (Assaf and Basser, 2005; Tuch et al., 2002; Yoshiura et al., 2001). Because of its Cartesian sampling, DSI may provide certain advantages over shelled schemes when estimating the EAP (Wedeen et al., 2008). Furthermore, measures derived from DSI acquisitions may be sensitive to complex biological changes like neuroinflammation, axonal loss and demyelination (Wang et al., 2021, 2020; Yeh et al., 2016, 2013; Zhang et al., 2013). Together, available research suggests immense potential for DSI. However, DSI’s Cartesian sampling scheme comes at the cost of a much longer acquisition time: a typical DSI scan can take upwards of twenty minutes to acquire, limiting its applications in both time-constrained research settings and clinical practice.

One potential solution to this obstacle is compressed sensing (CS) (Candes and Wakin, 2008; Donoho, 2006). CS is a widely used technique that can reconstruct signals from under-sampled data, and has been very valuable for advances in diverse fields including telecommunications (Berger et al., 2010), astronomy (Bobin and Starck, 2009; Wiaux et al., 2009), radar (Baraniuk and Steeghs, 2007; Herman and Strohmer, 2009), and medical imaging (Carroll et al., 2009; Lustig et al., 2007). In MRI, CS is often used to accelerate acquisitions by under-sampling *k*-space (Adluru and Dibella, 2008; He et al., 2007; Jaspan et al., 2015; Lustig et al., 2008). However, recent work has shown that dMRI acquisitions can under-sample *q*-space and still accurately estimate EAPs and other derivatives after using CS methods. (Menzel et al., 2011b; Merlet and Deriche, 2013; Michailovich et al., 2011; Paquette et al., 2015; Pu et al., 2011; Ramirez-Manzanares et al., 2007). Prior work has provided some guidance regarding optimal *q*-space sampling strategies (Menzel et al., 2011b; Merlet and Deriche, 2013) and effective signal recovery methods (Bilgic et al., 2012; Cheng et al., 2011; Gramfort et al., 2012; Tristán-Vega and Westin, 2011; Ye et al., 2012). These sampling and recovery methods have been validated in simulated data (Menzel et al., 2011b; Merlet and Deriche, 2013), post-mortem tissue (Jones et al., 2021), and in small samples of *in vivo* human data (n<3 (Bilgic et al., 2013, 2012; Merlet and Deriche, 2013)). While these studies have underscored CS-DSI’s promise, most of this work has primarily focused on sequence development and reconstruction techniques and no study to our knowledge has compared CS-DSI outputs to full DSI outputs in living humans. As such, important benchmarks of *in vivo* validity, including the accuracy and reliability of commonly used derivatives such as white matter bundles and scalar maps, have not been established. Furthermore, as the scan time for CS-DSI schemes is directly proportional to the degree to which the *q*-space is under-sampled, the trade-off between under-sampling and robustness remains unknown.

We sought to establish the validity of CS-DSI in living humans. We were motivated by the possibility that CS-DSI could provide accurate measures of white matter anatomy in a fraction of the scan time required by a full DSI scan. Such an advance would allow both researchers and clinicians to harness the advantages of DSI sequences that were previously impractical to deploy. We examined six different CS-DSI schemes and evaluated whether these schemes could generate white matter bundles and scalar maps comparable to those generated by a full DSI scheme. While there are many ways to sparsely sample *q*-space, we focused on the homogenous angular sampling scheme (HA-SC) (Merlet, 2013). HA-SC schemes ensure homogenous angular coverage of *q*-space and have been shown to produce more consistent reconstructions than other sampling strategies (Jones et al., 1999; Menzel et al., 2011b). To assess the relationship between CS-DSI validity and scan time, we included a range of sampling densities in our schemes that provided between 50-80% reduction in scan time. We capitalized on a unique dataset of twenty-six participants who were scanned over eight independent sessions using a full DSI scheme. From this full DSI scheme, we subsampled images to retrospectively generate synthetic CS-DSI images. Using these images, we compared CS-DSI to full DSI on the accuracy and test-retest reliability of commonly used derivatives. Finally, we replicated our results in an independent dataset of 20 participants who were prospectively scanned on a different scanner using CS-DSI acquisition schemes. As described below, we found that CS-DSI schemes offer highly accurate and reliable alternatives to the full DSI scheme, while only requiring a fraction of the scan time.

## 2 Methods

### 2.1 Overview

Our full DSI scheme sampled 258 points on a Cartesian grid in *q*-space. The CS-DSI schemes we evaluate here (**Figure 1**) are all subsets of this full DSI scan. As such, we can simulate their acquisition retrospectively by extracting the volumes contained in each CS sampling scheme from a full DSI acquisition. The dataset of full DSI acquisitions included 26 participants scanned over 8 sessions (average time between sessions = 14 days), acquiring the same full 258-direction scheme at each scan session (Cieslak et al., 2018; Nakuci et al., 2022). We then sub-sampled six different CS-DSI schemes from this full DSI dataset. Four of these schemes (HA-SC92, HA-SC55-1, HA-SC55-2, RAND57) were designed such that concatenating them would result in the full DSI scheme. To explore a larger range in the number of directions sampled, we generated two additional CS-DSI schemes (HA-SC92+55-1, HA-SC92+55-1) by combining pairs of these sub-sampled CS-DSI schemes (**Figure 1**). To additionally assess whether the behavior of retrospectively assembled CS-DSI schemes could be replicated in new data, we prospectively acquired four of the CS-DSI schemes that together made up the full 258-direction grid on an additional 20 participants who were each scanned once (**Figure 2**). For all of the CS-DSI schemes, we first extrapolated a full DSI image using an iterative L2-regularized algorithm (Bilgic et al., 2012). For brevity, we will refer to this extrapolated full DSI image as the CS-DSI image. These CS-DSI images were then reconstructed with Generalized Q-Sampling Imaging (GQI) in both datasets (Yeh et al., 2010).

**Figure 1:**
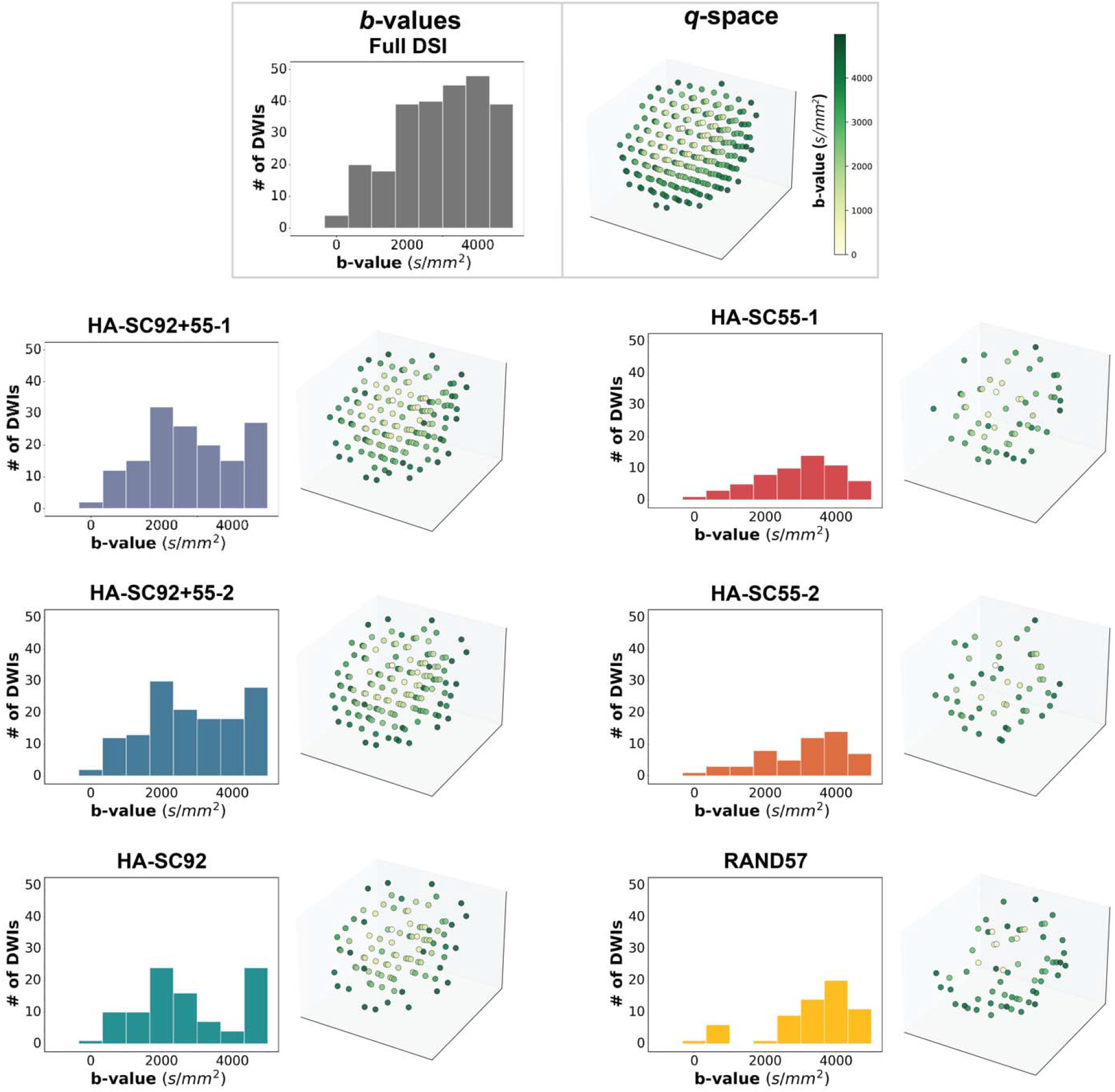
Description of the examined diffusion schemes with the histogram of *b*-values and distribution of *b*-vectors in *q*-space. *B*-vectors are scaled, and color coded by *b*-value. HA-SC: Homogenous Angular Sampling Scheme; RAND: Random Sampling Scheme.

**Figure 2:**
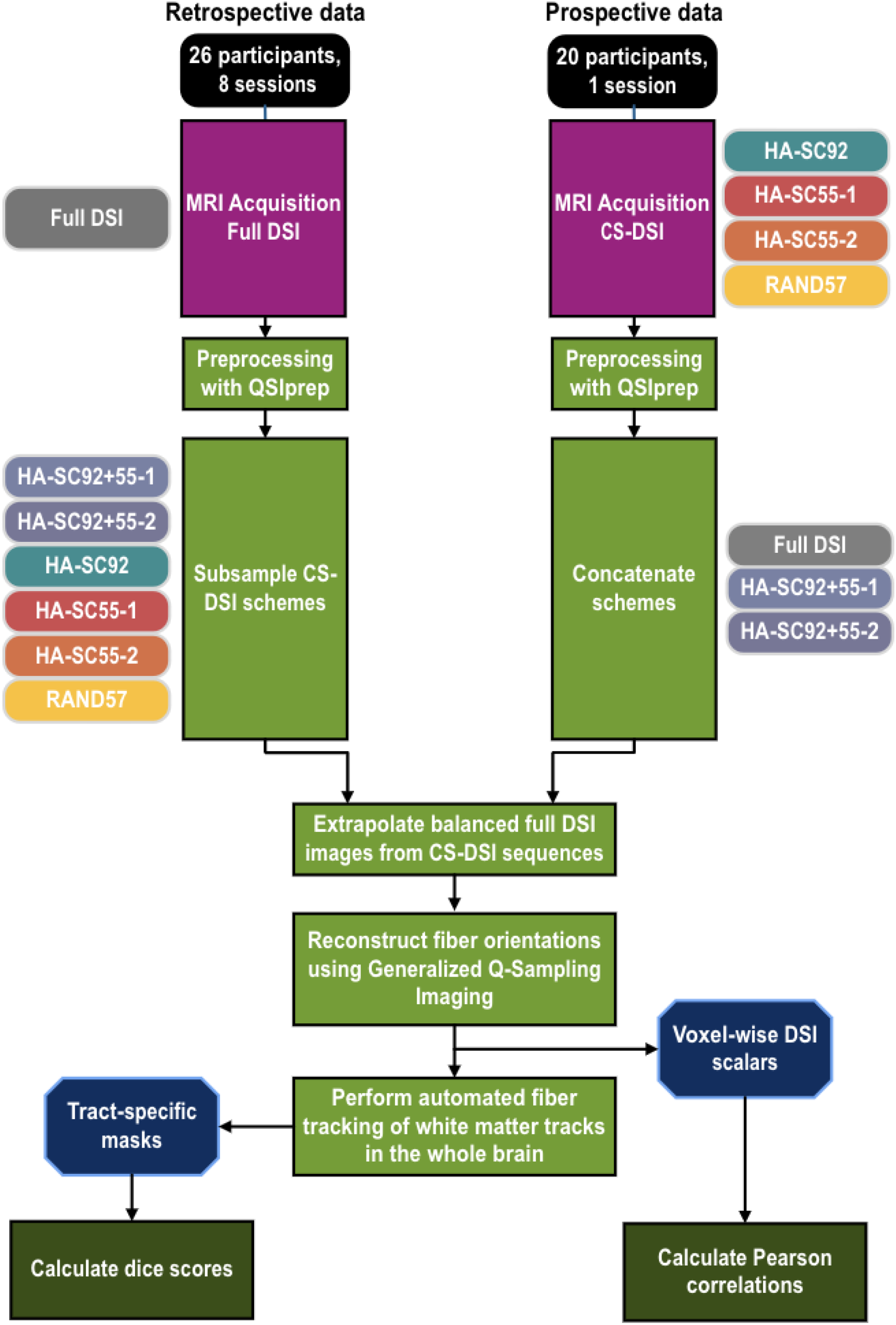
Preprocessing schematic and analytic overview.

We evaluated the CS-DSI image’s ability to segment bundles and generate voxel-wise scalars on two criteria: accuracy and reliability. We defined accuracy as the similarity between a measure derived from a CS-DSI image and that derived from the full DSI image (the acquired image for the retrospective data, and the concatenated image for the prospective data). As described below, we evaluated accuracy both within- and across scan sessions in the multi-session retrospective data; only same-scan accuracy could be evaluated in the prospective data. Inter-scan reliability was operationalized as the similarity between a given measure derived from the same diffusion scheme in different scan sessions; this could only be calculated for the multi-session retrospective data. Throughout, we compared bundles using calculated Dice scores and scalar maps using Pearson correlations.

### 2.2 Data Acquisition

#### 2.2.1 Retrospective Full DSI Data

The retrospective imaging data was extracted from a repeated measures dataset containing 8 full DSI scans (average time between sessions = 14 days) in a group of 26 healthy adults (mean age 22 ± 3.5 years, 16 Female) (Cieslak et al., 2018; Nakuci et al., 2022). CS-DSI schemes were created by extracting volumes from the full DSI scan. The participants were scanned using a Siemens Prisma 3T MRI scanner. Fitted padding was used to minimize head movements. Each dMRI scan was acquired with a full DSI scheme (i.e., Q5, half-sphere) comprised of 258 images with variable *b*-values (b_max_□=□5000 s/mm^2^) corresponding to 258 different isotropic grid points in *q*-space. Bipolar diffusion encoding gradients were used. The in-plane resolution and slice thickness were both set to 2mm. The images were acquired with a TE of 100.2 ms and a TR of 4300 ms. The total scan time was 20 minutes (**Table 1**). Due to incomplete data, two participants were removed from the final dataset, giving a total of 24 participants. We refer to this dataset as the “retrospective” dataset.

**Table 1:** Acquisition time for each scheme. * These schemes were concatenated from acquired schemes.

| Scheme | Number of unique directions<br>(including b=0) | Acquisition time (minutes) |
| --- | --- | --- |
| Full DSI | 258 | 20 |
| HA-SC92+55 – 1* | 146 | 11.9 |
| HA-SC92+55 – 2* | 146 | 11.7 |
| HA-SC92 | 92 | 7.4 |
| HA-SC55 – 1 | 55 | 4.5 |
| HA-SC55 – 2 | 55 | 4.3 |
| RAND57 | 57 | 5.4 |

#### 2.2.2 Prospective CS-DSI Data

After initial analysis of retrospectively synthesized CS-DSI schemes, we sought to replicate our results with prospectively acquired CS-DSI. All participants provided informed consent before participation in this study. All experimental procedures were approved by the University of Pennsylvania Review Board. We scanned 20 healthy adults (mean age = 26.5 ± 3.5 years, 9 Female) using a Siemens Prisma 3T MRI scanner under four different Compressed Sensing (CS) acquisition schemes (**Figure 1**). Three of the four CS schemes were homogenous angular sampling schemes (HA-SC) (Paquette et al., 2015) sampling 92, 55, and 55 directions respectively. HA-SC schemes are sampling schemes that ensure a random but uniform angular covering of the *q*-space (*Figure 1)*. Note that some points in *q*-space were repeated across the three HA-SC schemes to ensure consistency in the big and little deltas across scans. The fourth scheme was a random scheme (RAND) sampling 57 directions. In the RAND scheme, samples were not uniformly distributed and were rather designed such that combining the RAND scheme with the unique samples of all the acquired HA-SC schemes resulted in the balanced full DSI scheme used in the retrospective dataset (**Figure 1**). The in-plane resolution and slice thickness were both set to 2mm with a multiband acceleration factor of 4. The volume images were acquired using a TE of 90 ms and a TR of 4300 ms. The acquisition times for the CS-DSI schemes were significantly lower than that for the full DSI scheme (*Table 1*). We refer to this dataset as the “prospective” dataset.

### 2.3 Preprocessing

Preprocessing was performed using *QSIPrep* 0.4.0 (Cieslak et al., 2021) which is based on *Nipype* 1.1.9 (Gorgolewski et al., 2011, 2017) on the retrospective data and *QSIPrep* 0.8.0 (*Nipype* 1.4.2) on the prospective data. *QSIprep* was run on the full DSI scan in the retrospective dataset and separately on each of the 4 directly acquired CS-DSI scans in the prospective dataset. For all full DSI and CS-DSI scans, initial motion correction was performed using only the b=0 volumes. An unbiased b=0 template was constructed over 3 iterations of affine registrations. The SHORELine method (Cieslak et al., 2022) was used to estimate head motion in *b*>0 volumes. SHORELine is a cross-validated method that entails leaving out each *b*>0 image and fitting the 3dSHORE basis function (Ozarslan et al., 2013) to all the other volumes using L2-regularization. The signal for the left-out volume serves as the registration target. A total of 2 iterations were run using an affine transform. Model-generated volumes were transformed into alignment with each *b*>0 volume. Both slice-wise and whole-brain quality control measures (cross correlation and R^2^) were calculated. No susceptibility distortion correction was performed. The DWI time-series were resampled such that they were aligned to anterior and posterior commissures (ACPC), generating a preprocessed DWI run in ACPC space. The accuracy of *b*-table orientation was examined by comparing fiber orientations with those of a population-averaged template (Schilling et al., 2019).

The four CS schemes used in acquiring the prospective data were designed such that combining them resulted in a full DSI image. To assess whether CS schemes longer than the ones acquired could generate results closer to the full DSI, two additional CS schemes were also generated by combining the preprocessed HA-SC92 scheme with each of the preprocessed HA-SC55 schemes (forming HA-SC92+55-1 and HA-SC92+55-2). For each participant in the retrospective data, the 6 CS-DSI schemes mentioned above were generated from the full-DSI scheme by extracting out only the relevant volumes from the full preprocessed image. This distinction is important to note as the object of comparison for CS-DSI derivatives are their corresponding full-DSI derivatives. Critically, in the retrospective data, the full-DSI is acquired in a single scan where all images are preprocessed together. The prospective full-DSI is composed of four individual scans that were each preprocessed separately and then concatenated.

### 2.4 CS Reconstruction

Following preprocessing, we used DSI Studio to perform fiber tracking and to derive scalar metrics. GQI requires balanced q-space sampling, so a full, balanced 258-direction DSI image was extrapolated from the CS-DSI schemes. Extrapolation was performed via CS using the 3DSHORE basis function (Ozarslan et al., 2013) with a radial order of 8 and L2 regularization. For brevity, we will refer to this extrapolated full DSI image as the CS-DSI image from now on. These GQI reconstructions with a diffusion sampling length ratio of 1.25 were used as the input for fiber tracking and anisotropy scalar estimation (**Figure 2**).

### 2.5 Tractography and Bundle Segmentation

A deterministic fiber tracking algorithm (Yeh et al., 2013) was used with three augmented tracking strategies: parameter saturation, atlas-based track recognition, and topology-informed pruning (Yeh, 2020) to improve reproducibility. White matter bundles were segmented using a bundle atlas from Yeh et al., 2018 in the whole brain with a distance tolerance of 16.00 (mm) in the ICBM152 space. The track-to-voxel ratio was set to 2. Streamlines with lengths shorter than 30 mm or longer than 300 mm were discarded. Topology-informed pruning (Yeh et al., 2019) was applied to the tractography with 32 iteration(s) to remove false connections. Shape analysis (Yeh, 2020) was conducted to derive shape metrics for tractography. A total of 56 fiber bundles were segmented in this way (**Supplementary Table 1**). Each bundle was segmented 1344 times for the retrospective dataset (24 participants, 8 scan sessions, 7 diffusion schemes) and 140 times for the prospective dataset (20 participants, 1 session, 7 diffusion schemes). Note however that the following bundles could not be segmented for some of the schemes in some of the participants: Left Vertical Occipital Fasciculus (507 failures), Right Vertical Occipital Fasciculus (250 failures), Left Parahippocampal Cingulum (118 failures), Right Parahippocampal Cingulum (153 failures), and the Left Fornix (1 failure). Failures were observed evenly across all schemes and were excluded from the final analysis.

The segmented bundles were then converted into binary masks, and Dice scores were calculated between the masks for each scenario using the following formula:

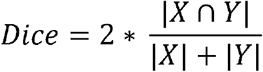

High Dice scores denoted high overlap between the two bundles examined and vice-versa (**Figure 3**). For each CS scheme and bundle, accuracy of bundle segmentation was defined as the Dice score between the bundle generated by the CS image and the bundle generated by the corresponding full DSI image. Two different types of accuracies were calculated: same-scan accuracy and inter-scan accuracy. Same-scan accuracy was defined as the Dice score between the bundle generated by the CS-DSI image and the corresponding bundle generated by the full DSI image (the acquired full image for the retrospective data, and the combined image for the prospective data) for a participant, *within the same scan session*. Hence each participant in the retrospective dataset had 8 values for same-scan accuracy (one for each scan session) for a given bundle and diffusion scheme, and participants in the prospective dataset had one value. To account for the noise between scan sessions, inter-scan accuracy was defined as the Dice score between the bundle generated by the CS-DSI image in one scan session and the bundle generated by the full DSI image in *another scan session*, for the same participant. Inter-scan accuracy was calculated for each pair of scan sessions, so each participant in the retrospective dataset had 56 values for inter-scan accuracy (one for each permuted pair of scan sessions).

**Figure 3:**
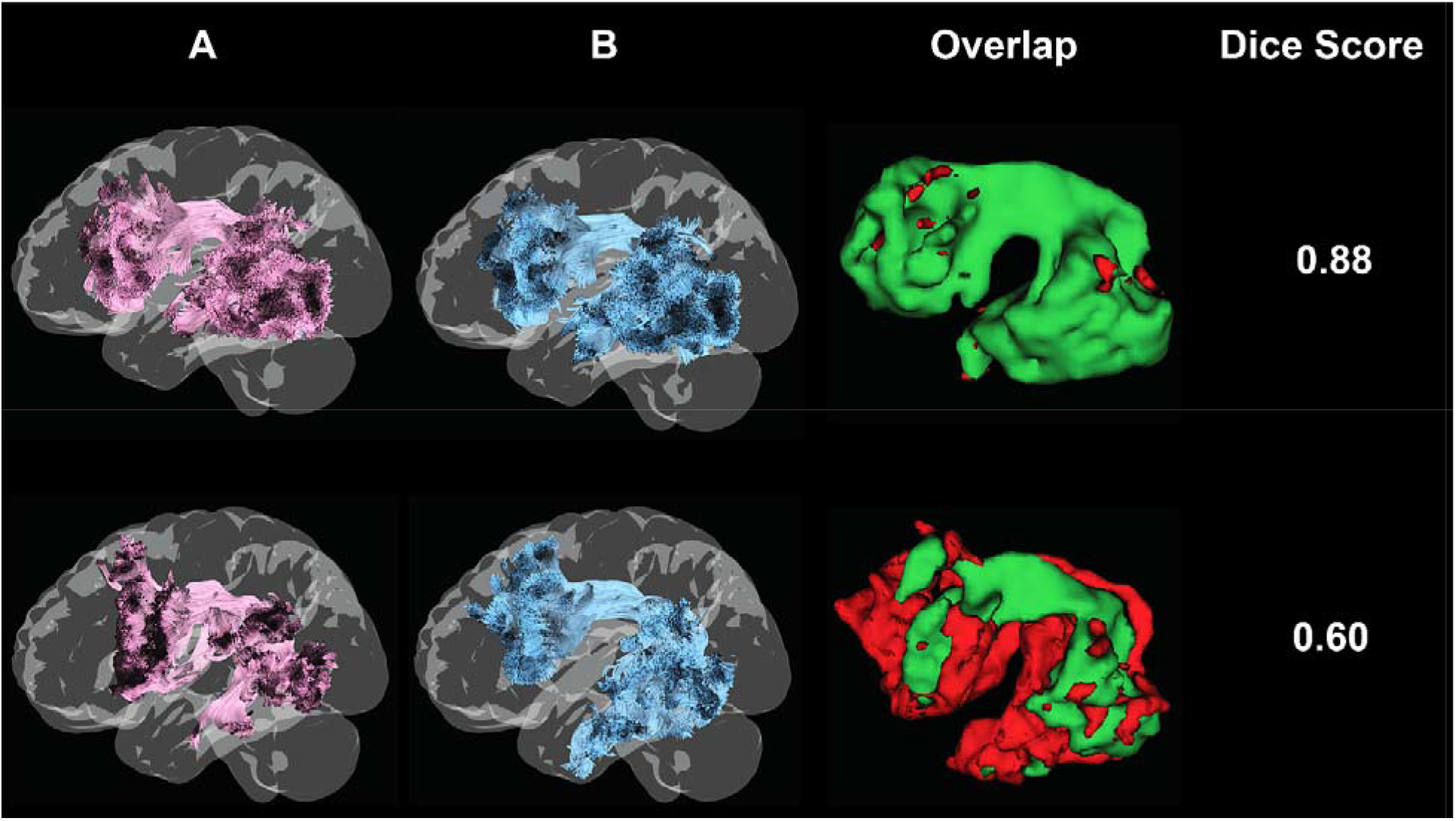
Graphical representation of range of Dice scores. The exemplar bundle shown here is the left Arcuate Fasciculus. The first two columns (A and B) show the left Arcuate Fasciculus derived from different DSI images (Top row: same participant, different full DSI sessions; Bottom row: same participant and session, RAND57 and full DSI). The third column shows voxels that overlap between the bundles (green), and differences (red).

Similarly, inter-scan reliability for a given diffusion scheme and bundle was defined as the set of Dice scores between the bundles generated by each pair of scan sessions. Notably, inter-scan accuracy compared pairwise session differences <u>between</u> the CS-DSI and full DSI images, while inter-scan reliability compared pairwise session differences <u>within</u> a scheme. Inter-scan reliability was also calculated independently for each participant in the retrospective dataset. As above, given the number of session pairs, each participant had 28 values for reliability (one for each unique pair of scan sessions) for a given bundle and diffusion scheme (**Table 2**). Inter-scan metrics could not be calculated in the prospective dataset because of the absence of multi-session data.

**Table 2:** Definitions of validity metrics used to compare diffusion schemes. The relationship is defined by the Dice score when the derivatives are bundle segmentations, and by the Pearson correlation when the derivatives are voxel-wise scalars.

| Metric | Definition | Scheme | Datasets |
| --- | --- | --- | --- |
| Same-scan accuracy | Relationship between derivative produced by a CS-DSI image and the corresponding derivative produced by the full DSI image, within a participant and session | CS-DSI only | Both retrospective and prospective |
| Inter-scan accuracy | Relationship between derivative produced by a CS-DSI image in one session, and the derivative produced by the full DSI image in another session, within a participant. Calculated between each pair of sessions. | CS-DSI only | Only retrospective |
| Inter-scan reliability | Relationship between derivative produced by a DSI image in one session, and the derivative produced by the same DSI image in another session, within a participant. Calculated between each pair of sessions. | CS-DSI and full DSI | Only retrospective |

### 2.6 Deriving whole-brain voxel-wise maps of white matter microstructure

While macrostructural derivatives like bundles are very useful for studying white matter connectivity, sometimes more localized measures are useful. Voxel-wise scalar metrics can provide additional insight into white matter structure and health and could capture pathology or other information at the microstructural level. GQI also allowed us to derive voxel-wise scalar maps, permitting more fine-grained comparisons (Yeh et al., 2010). Note that traditional tensor metrics cannot be used with these data as tensor assumptions may not hold true for higher *b*-values where diffusion is non-Gaussian. Accordingly, we examined three DSI-based scalar metrics that quantified different fiber properties: Normalized Quantitative Anisotropy (NQA), Generalized Fractional Anisotropy (GFA) and the Isotropic value (ISO) (Yeh et al., 2013, **Figure 4**). All three metrics were calculated from the orientation distribution function (ODF) of each voxel. Quantitative anisotropy (QA) is the value of the highest peak of the ODF. QA is similar to the Fractional Anisotropy (FA) measure derived from tensors, except QA is defined per fiber population and so is less affected by crossing fibers. QA has arbitrary units, so we scaled the maximum QA of a subject to 1 to generate the more easily comparable Normalized QA (NQA). In contrast to NQA, GFA is defined as the standard deviation between all vertices of the ODF divided by the root mean square of the vertices. Both GFA and NQA are measures of directional integrity, but GFA is more sensitive to differences in ODF shapes, as broadly different ODF structures can have the same maxima but differing peak variances. Finally, ISO is the minima of the ODF, and mainly measures background isotropic diffusion from CSF (Yeh et al., 2013). Together, these scalars describe diverse properties of a given voxel. For each scalar metric examined, we calculated the Pearson correlation coefficient between whole-brain maps generated by CS-DSI and full DSI images. As with the Dice scores, the same-scan accuracy of each metric was defined as the Pearson correlation between the voxel-wise metric calculated from the CS-DSI image and that calculated from the full DSI image within a particular scan session. The inter-scan accuracy was the Pearson correlation between the voxel-wise metric calculated from the CS-DSI image in one scan session and that calculated by the full DSI image in a different scan session, for each pair of sessions. The inter-scan reliability was the set of Pearson correlations between voxel-wise metrics derived from each pair of scan sessions within a single diffusion scheme (see **Table 2**).

**Figure 4:**
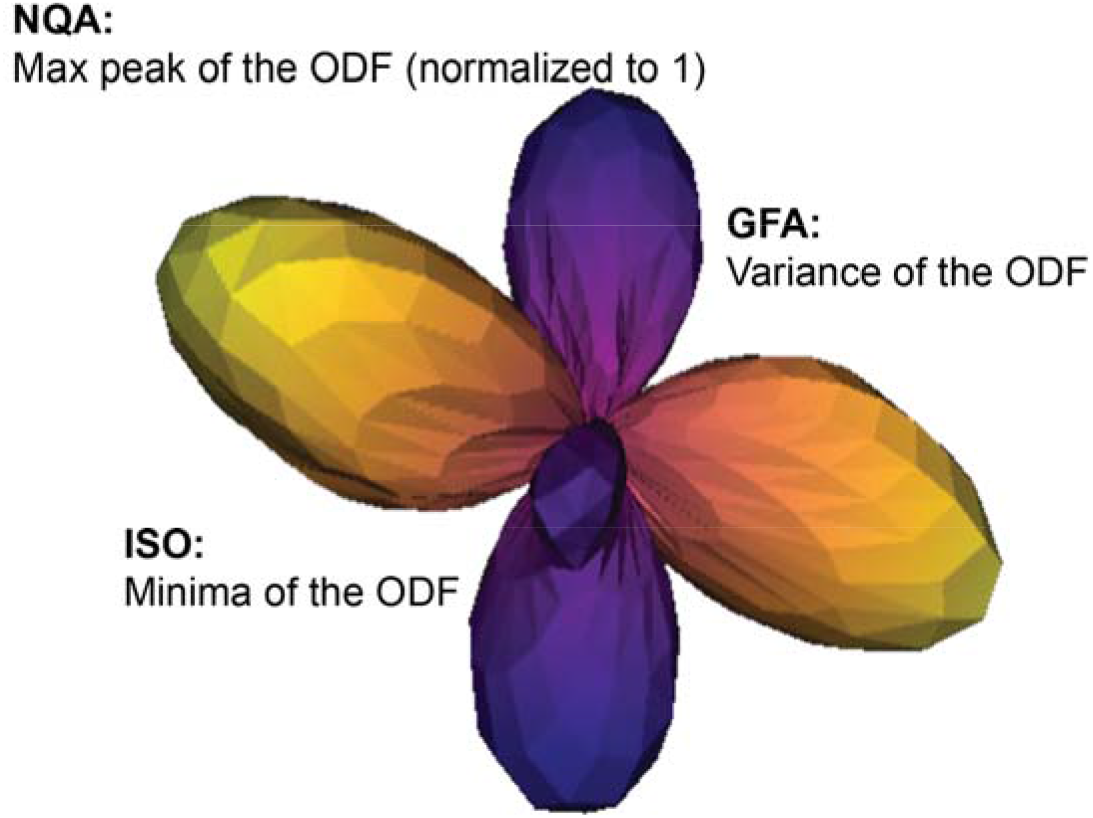
Schematic of DSI scalars examined.

### 2.7 Statistical Analysis

Each of the validity metrics described in **Table 2** were calculated for every participant, session, and bundle/scalar. The inter-scan reliability of the full DSI (from the retrospective dataset) was used as the “gold” standard for all comparisons. Therefore, for each bundle/scalar, the distributions of each of the validity metrics of CS-DSI across all participants were compared with the distribution of the inter-scan reliability of the full DSI. To determine if there was a statistically significant difference between these distributions, we compared the median difference between these distributions against a null distribution generated by permutation testing (see **Supplementary**). Two-sided *p*-values were calculated across subjects per bundle for the Dice score comparisons. We corrected for multiple comparisons (56 bundles for the Dice scores, and 3 metrics examined for the scalars) using the Benjamini/Hochberg false discovery rate method at 5% (Benjamini and Hochberg, 1995).

## 3 Results

### 3.1 CS-DSI schemes can segment major fiber bundles

One of the main applications of dMRI is to delineate white matter bundles. We first qualitatively examined whether both retrospectively generated and prospectively acquired CS-DSI images could segment white matter bundles of varying shapes, convergent with bundles segmented by the corresponding full DSI image (i.e., the acquired full DSI image for the retrospective data, and the concatenated full DSI image for the prospective data). Inspection of data from individual participants revealed that we could adequately segment bundles from all CS-DSI images (see exemplar results in **Figure 5**).

**Figure 5:**
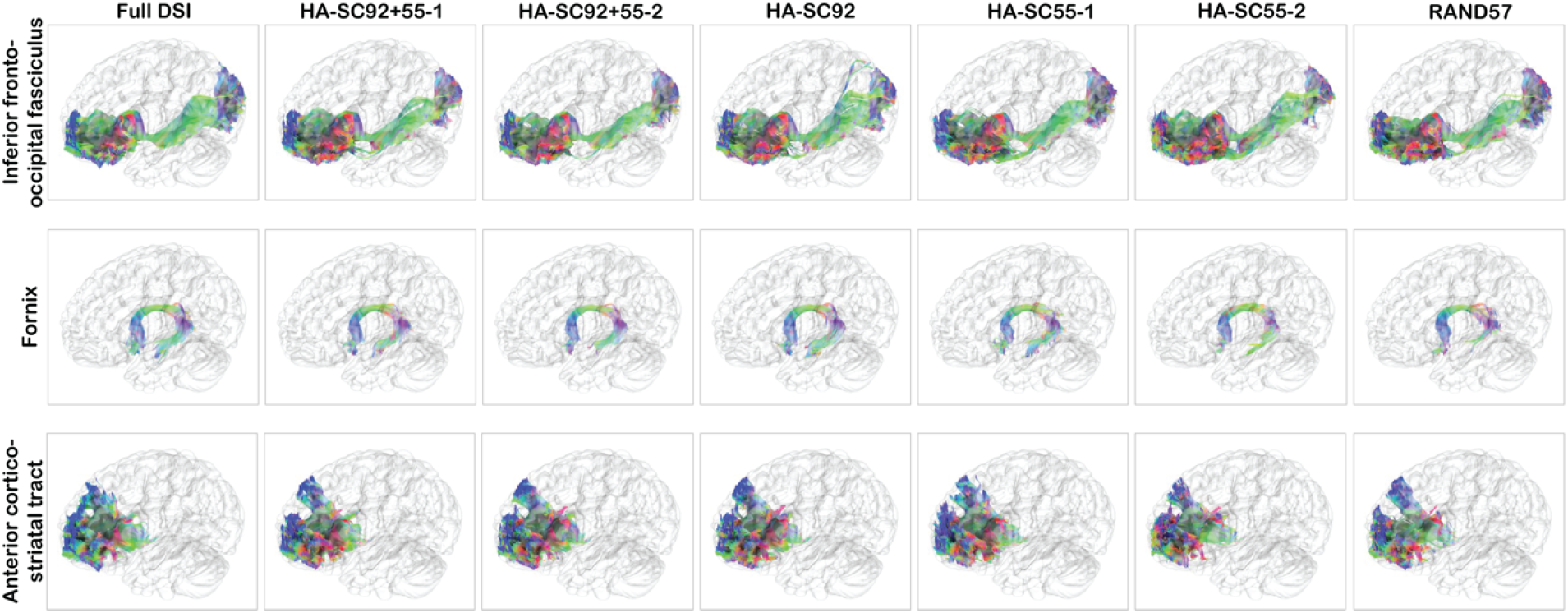
All CS images can segment both long-range and short-range bundles, comparable with those derived from a full DSI image. These figures are exemplar segmentations of different bundles in a single participant from the retrospective dataset.

Next, we quantitatively validated CS-DSI’s ability to delineate bundles and calculated voxel-wise scalars, comparing CS-derived results to those derived from the full DSI image. The inter-scan reliability of the full DSI image was selected as the gold standard for comparison, as it captured noise and other session-related differences. Inter-scan reliability was calculated from the retrospective dataset and was defined as the set of relationships between derivatives produced by each pair of sessions within a given diffusion scheme (**Table 2**).

### 3.2 Bundles segmented by CS-DSI images are similar to those segmented by their corresponding full DSI images in the same scan session

To quantitatively compare the bundles segmented by CS-DSI images and those segmented by full DSI images, we examined two measures of accuracy in the retrospective data: same-scan and inter-scan accuracy. Same-scan accuracy was defined as the Dice score between each bundle segmented by a CS-DSI image and the bundle segmented by the full DSI image for a given participant *within the same scan session* (**Table 2**). We found that all CS-DSI images showed high same-scan accuracies across fiber bundles (with median Dice scores ranging 0.72-0.82, **Figure 6a**). For additional context, we used full DSI inter-scan reliability as the standard of reference as test-retest differences are inevitable in neuroimaging, even when using longer full-grid schemes. We asked whether the difference between CS-DSI and full DSI bundles were comparable with test-retest differences of full DSI bundles. To this end, we compared the distributions of these same-scan accuracies of each CS-DSI scheme with the distributions of the full DSI inter-scan reliabilities across participants and sessions for every bundle generated. *P*-values were calculated per bundle using permutation testing on the median differences between these distributions (see **Supplementary**). For the more densely sampled CS-DSI schemes, like HA-SC92+55-1, HA-SC92+55-2 and HA-SC92, the same-scan accuracy was statistically the same as the full DSI inter-scan reliability (FDR-corrected *p*-values > 0.1 for all bundles where full DSI reliability > CS-DSI accuracy, **Supplementary Table 2**). As expected, the schemes with fewer directions sampled had an overall lower segmentation accuracy, and the RAND57 scheme performed worse than all the uniformly sampled HA-SC schemes. While the sparser sampling schemes showed significant differences across almost all bundles, the median difference between CS-DSI same-scan accuracy and full DSI inter-scan reliability was still quite small (HA-SC55-1 = 0.039, HA-SC55-2 = 0.058, RAND57 = 0.074, **Figure 6a, Supplementary Table 2)**. These results suggest that bundles from CS-DSI are comparable to bundles generated using the full DSI grid scheme, and that accuracy scales with the degree of under-sampling.

**Figure 6.**
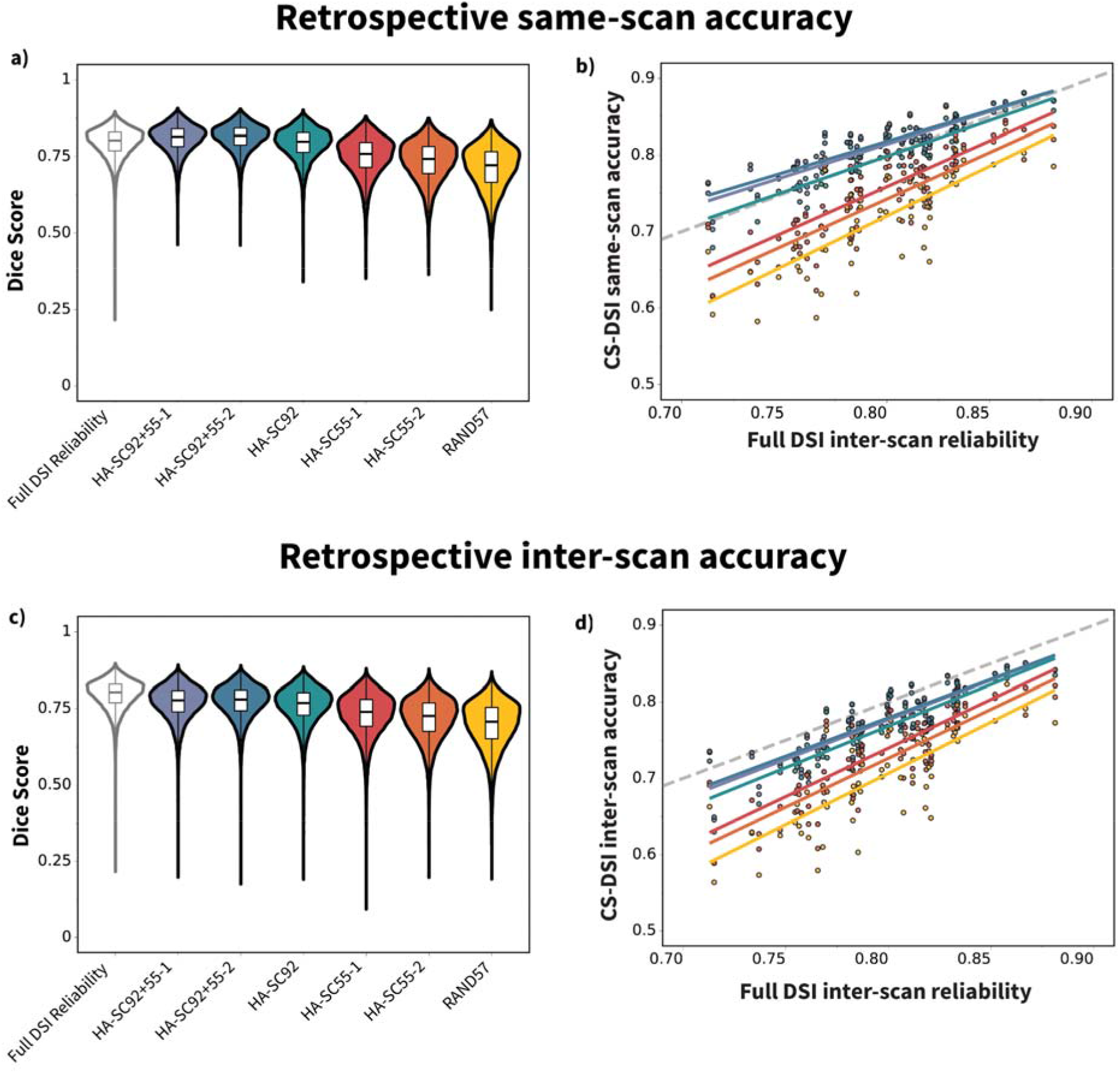
CS-DSI bundle segmentation accuracy is comparable and correlated with full DSI inter-scan reliability both within and across scan sessions. **a)** Same-scan accuracy of CS-DSI schemes is comparable with inter-scan reliability of the full DSI scheme. Violins represent distributions of Dice scores across all bundles. **b)** Median distribution of the same-scan accuracy for each CS-DSI scheme is highly correlated with the median distribution of the inter-scan reliability of the full DSI scheme. Here, each point on the scatter plot represents the median of Dice scores for a single bundle across participants and sessions. The gray dashed line denotes x=y. **c)** Inter-scan accuracy of CS-DSI schemes is comparable with inter-scan reliability of the full DSI scheme. Violins represent distributions of Dice scores across all bundles. **d)** Median distribution of the inter-scan accuracy for each CS-DSI scheme is highly correlated with the median distribution of the inter-scan reliability of the full DSI scheme. Here, each point on the scatter plot represents the median of Dice scores for a single bundle across participants and session pairs. The gray dashed line denotes x=y.

Not all bundles are reliably segmented, even when a full DSI image is used for tractography (**Supplementary Table 3**). As such, we next asked whether the accuracy of CS-based bundle segmentations was related to how reliably that bundle was segmented with a full DSI scheme. For example, the left and right reticular tracts had low same-scan accuracy across most CS-DSI schemes. We reasoned that this might be due to poorer segmentation reliability of this bundle even when using full DSI. Across bundles, we found that CS-DSI same-scan accuracy was strongly correlated with full DSI inter-scan reliability (Pearson R ranging from 0.78-0.89; *p* < 0.001 for all CS-DSI schemes, **Figure 6b**). This suggests that bundles that are not reliably segmented by the full DSI images were also less likely to be accurately segmented by any of the CS-DSI images. Moreover, the more sparsely sampled CS-DSI images were more severely affected by this relationship and yielded lower Dice scores in bundles that showed low inter-scan reliability with full DSI. Bundles with high inter-scan reliability with full DSI were more accurately segmented by all CS-DSI images, including the more sparsely sampled schemes. Notably, the HA-SC92+55-1 and HA-SC92+55-2 schemes had same-scan accuracies that were *higher* than the inter-scan reliability of the full DSI for most bundles (**Figure 6b**). This is likely due to the fact that differences between scan sessions were greater than differences introduced by subsampling of the same input data.

### 3.3 Bundles segmented by CS-DSI images are also similar to those segmented by full DSI images acquired in a different scan session

The prior analysis compared bundles from the full DSI images to bundles generated from sub-sampled CS-DSI images within the same scan. To further examine CS-DSI segmentation accuracy while accounting for scan differences, we examined a second measure: inter-scan accuracy (**Table 2**). We defined inter-scan accuracy as the set of Dice scores between CS-DSI bundles and full DSI bundles in disparate sessions. Understandably, the inter-scan accuracies were lower than the same-scan accuracies on average, as they were derived from different sessions (median Dice scores ranging from 0.71-0.78, **Supplementary Table 4**). This accuracy scaled with the number and uniformity of directions sampled, with dense HA-SC schemes performing with higher accuracy than the RAND scheme. While all bundles showed a significant difference between CS-DSI inter-scan accuracy and full DSI reliability for almost all diffusion schemes, the median difference between these distributions was very low (<0.09, **Figure 6c, Supplementary Table 4**). All CS-DSI schemes could still delineate very similar bundles with comparable anatomy to those delineated by full DSI. These results demonstrate that CS-DSI schemes can generate bundles similar to those segmented by the full DSI scheme in a different scan session.

We next asked if the CS-DSI inter-scan accuracy of specific bundles similarly scaled with their full DSI inter-scan reliability. We again found that bundles that were more reliably segmented by full DSI had higher inter-scan accuracies, and vice-versa (Pearson R ranging from 0.78-0.88; *p*<0.001, **Figure 6d)**. As observed previously, the slopes of this relationship scaled with sampling density, and the more sparsely sampled schemes were more severely affected by low inter-scan reliability in the full DSI scheme. These results emphasize that most bundles can be segmented by CS-DSI schemes with very high accuracies.

### 3.4 The inter-scan reliability of bundles from CS-DSI is comparable to full DSI

Having demonstrated that CS-DSI schemes could segment bundles accurately, we next asked whether the CS-DSI segmented bundles were as reliable between scan sessions as bundles generated from the full DSI scheme. For each diffusion scheme, we calculated the inter-scan reliability for each bundle as the set of Dice scores between each pair of scan sessions within a participant (**Table 2**). We found that the inter-scan reliabilities across bundles were high for all CS-DSI schemes (medians ranging from 0.73-0.78). We then compared the distributions of inter-scan reliabilities for each CS-DSI scheme to the full DSI scheme across all participants within each bundle. We again found that the median difference between these distributions scaled with sampling density (**Figure 7a**). However, these median differences were low (<0.07, **Supplementary Table** *5*) and after correcting for multiple comparisons, no bundle showed a significant difference between CS-DSI and full DSI reliabilities for any scheme (*p* > 0.1, **Supplementary Table 5)**. These results demonstrate that CS-DSI is as reliable as full DSI for segmenting bundles.

**Figure 7:**
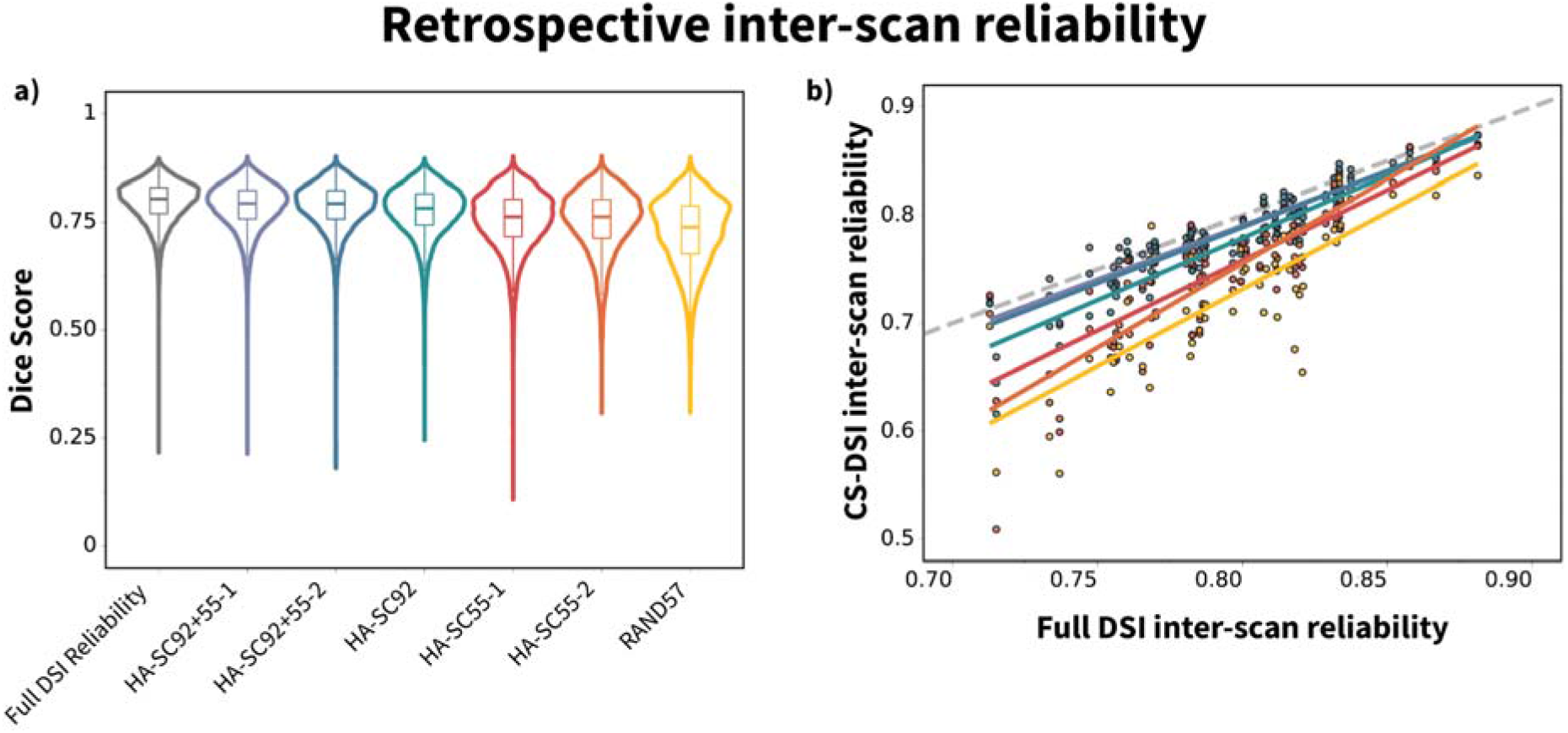
CS-DSI inter-scan reliability of bundle segmentation is comparable to and correlated with full DSI inter-scan reliability. **a**) CS-DSI inter-scan reliability is comparable with full DSI inter-scan reliability. Violins represent distributions of Dice score across bundles and participants. **b)** Median distribution of the inter-scan reliability for each CS-DSI scheme is highly correlated with the median distribution of the inter-scan reliability of the full DSI scheme. Here, each scatter dot represents the across-participant median of Dice scores for a single bundle. The gray dashed line denotes x=y.

Mirroring what we observed for bundle segmentation accuracy, we found that the inter-scan reliability of the Dice scores for all CS-DSI schemes was correlated with the inter-scan reliability of the full DSI scheme (Pearson R ranging from 0.80-0.94; *p < 0*.*001*) **Figure 7b**). Bundles that were most susceptible to noise between sessions in the full DSI segmentations were also less reliably segmented in the CS-DSI images over sessions. Again, the inter-scan reliability of sparsely sampled CS-DSI schemes (like HA-SC55 and RAND57) was more impacted in bundles that showed lower inter-scan reliability in the full DSI scheme.

### 3.5 CS-DSI and full DSI produce highly correlated voxel-wise scalar maps in the same scan session

After showing that CS-DSI schemes studied could accurately and reliably segment white matter bundles, we next turned to voxel-wise scalar maps. We examined three different microstructural properties: NQA, GFA, and ISO (Yeh et al., 2013). We evaluated each of these voxel-wise scalars with the same validation metrics as done for bundle segmentation: same-scan accuracy, inter-scan accuracy, and inter-scan reliability (**Table 2**). We found that the CS-DSI derived voxel-wise scalars were all highly correlated with the full DSI derived voxel-wise scalars, demonstrating high same-scan accuracy. Moreover, the distribution of same-scan accuracies of all the CS-DSI maps was comparable with the distribution of inter-scan reliability of the full DSI maps for all the scalars studied (**Figure 8a-c**). For GFA and ISO, all HA-SC images generated scalars with no significant difference between their same-scan accuracy and the full DSI inter-scan reliability distributions (**Supplementary Table 6**). For NQA, there was no difference between the HA-SC55-1 image and full DSI. In contrast, HA-SC55-2 and RAND57 had significantly lower same-scan accuracies than full DSI inter-scan reliabilities, albeit with very low median differences (< 0.05).

**Figure 8:**
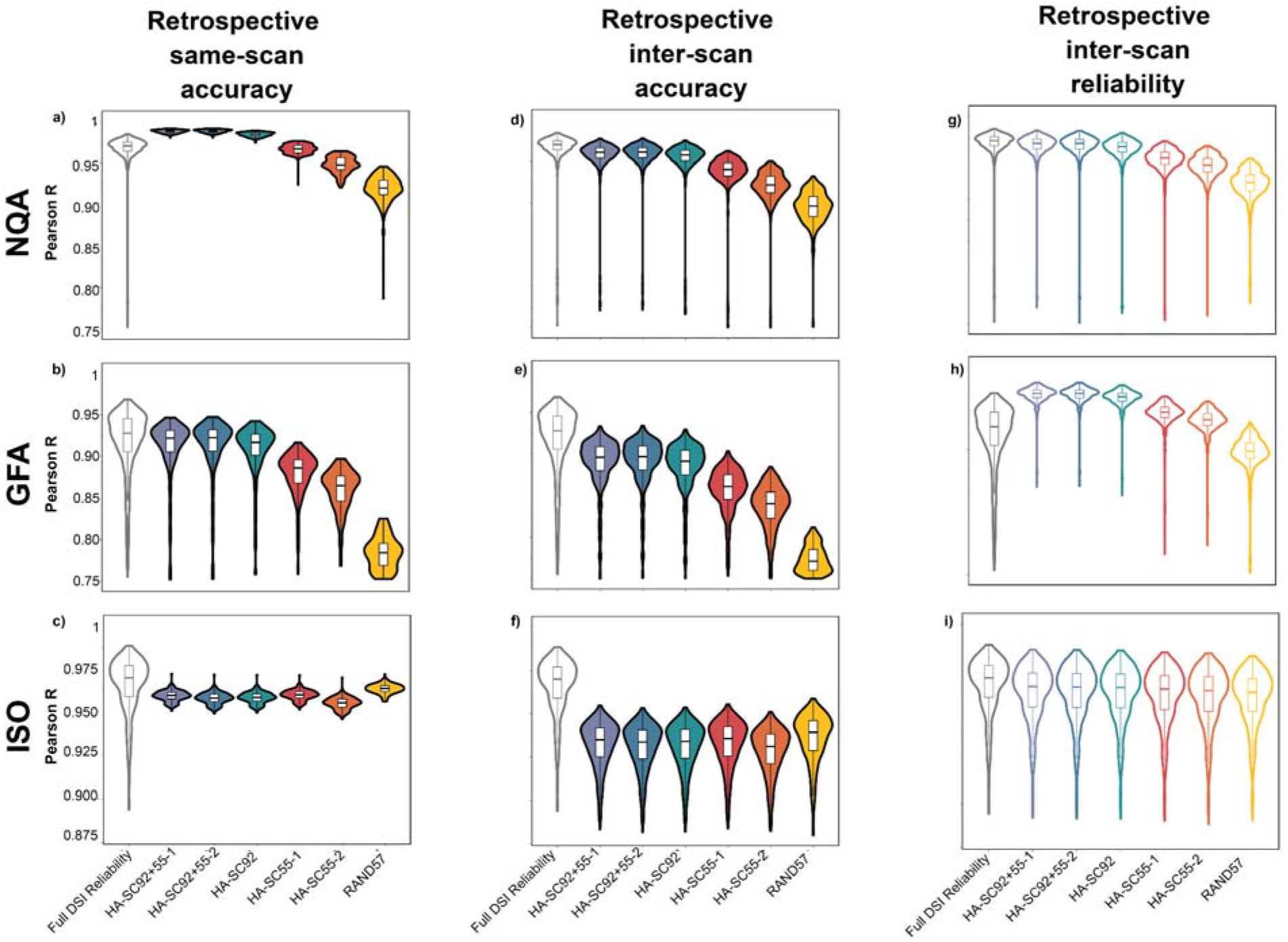
Comparing full DSI reliability with CS-DSI same-scan accuracy (a-c), inter-scan accuracy (d-f) and inter-scan reliability (g-i) when deriving whole-brain voxel-wise scalar maps.

Interestingly, HA-SC92+55-1, HA-SC92+55-2 and HA-SC92 had significantly higher same-scan accuracy for NQA as compared to the full DSI inter-scan reliability (**Figure 8a, Supplementary table 6**), which again is likely because the CS-DSI schemes and full DSI schemes within a scan session sub-sampled the same input data. Together, these results indicate that CS-DSI and full DSI produce similar estimates of voxel-wise scalar maps within the same scan session.

### 3.6 CS-DSI and full DSI produce similar estimates of voxel-wise scalar maps across scan sessions

As for the bundles, we next sought to further examine the accuracy of a scalar map of microstructure while controlling for the effect of scan session. Here, we defined inter-scan accuracy as the set of Pearson correlations between a CS-DSI derived voxel-wise scalar and the full DSI voxel-wise scalar in disparate sessions (**Table 2**). As expected, inter-scan accuracies were lower than the same-scan accuracies and scaled with the number and uniformity of directions sampled. These distributions were significantly different for all CS-DSI schemes and scalars examined (p < 0.05), but the median difference between the CS-DSI inter-scan accuracy and full DSI reliability was low for all HA-SC schemes (**Figure 8 d-f, Supplementary Table 7)**. Together these results demonstrate that CS-DSI schemes can generate whole brain voxel-wise scalars with high accuracy.

### 3.7 The inter-scan reliability of CS-DSI derived scalars is very similar to full DSI

Next, we examined how the inter-scan reliability of the voxel-wise scalar maps generated by the CS-DSI images compared with that of the full DSI image. As prior, we defined inter-scan reliability as the set of Pearson *R* values between a voxel-wise scalar metric generated by each pair of scan sessions. As for white matter bundles, we found that the ability of a CS-DSI image to reliably estimate any of the scalar metrics decreased with decrease in *q*-space sampling density (**Figure 8 g-i**). However, we found no significant differences in the inter-scan reliabilities between the CS-DSI derived scalars and the full DSI derived scalars (*p* > 0.05, **Supplementary table 8**). The median differences between these distributions were also very low across scalar metrics and diffusion schemes (<0.05). Interestingly, we found that all the HA-SC schemes had numerically (but not significantly) higher inter-scan reliabilities that the full DSI scheme for GFA, suggesting that strategic sub-sampling of *q*-space might be beneficial to reducing inter-scan noise in this metric. (**Figure 8h)**. These results suggest that CS-DSI schemes can be used to rapidly measure local microstructural properties with a similar reliability as a full DSI scheme.

### 3.8 CS-DSI accuracy can be replicated in an independent, prospectively acquired dataset

The results in the previous sections are from a dataset where synthetic CS-DSI data was generated *retrospectively:* full DSI schemes were collected and CS-DSI schemes were sub-sampled from the acquired data. Furthermore, because these CS-DSI images were subsampled *after* preprocessing the full DSI image, the CS-DSI volumes benefitted from better motion correction as they could rely on points in *q-*space that were not a part of the synthesized subsample. We next asked whether prospectively acquired CS-DSI data without this advantage could generate bundles and scalars at similar levels of accuracy. Note that in the prospective dataset, only a single session was acquired for each CS scheme, so we could not evaluate inter-scan accuracy or inter-scan reliability of the CS-DSI schemes. For this dataset, the full DSI image was constructed by concatenating the four acquired CS-DSI images (HA-SC92, HA-SC55-1, HA-SC55-2, RAND57). Same-scan accuracy was defined as the relationship between the CS-DSI derivative and the *concatenated* full DSI derivative for each scheme. Critically, we replicated the finding that all CS-DSI schemes had high same-scan accuracies across bundles (medians ranged from 0.74-0.80; **Figure 9a**). Furthermore, we replicated the result that bundle segmentation accuracy was highly correlated with the between-scan reliability, again using the full DSI reliability from the retrospective dataset as a reference (Pearson R ranging from 0.76-0.81, *p<0*.001, **Figure 9b**).

**Figure 9:**
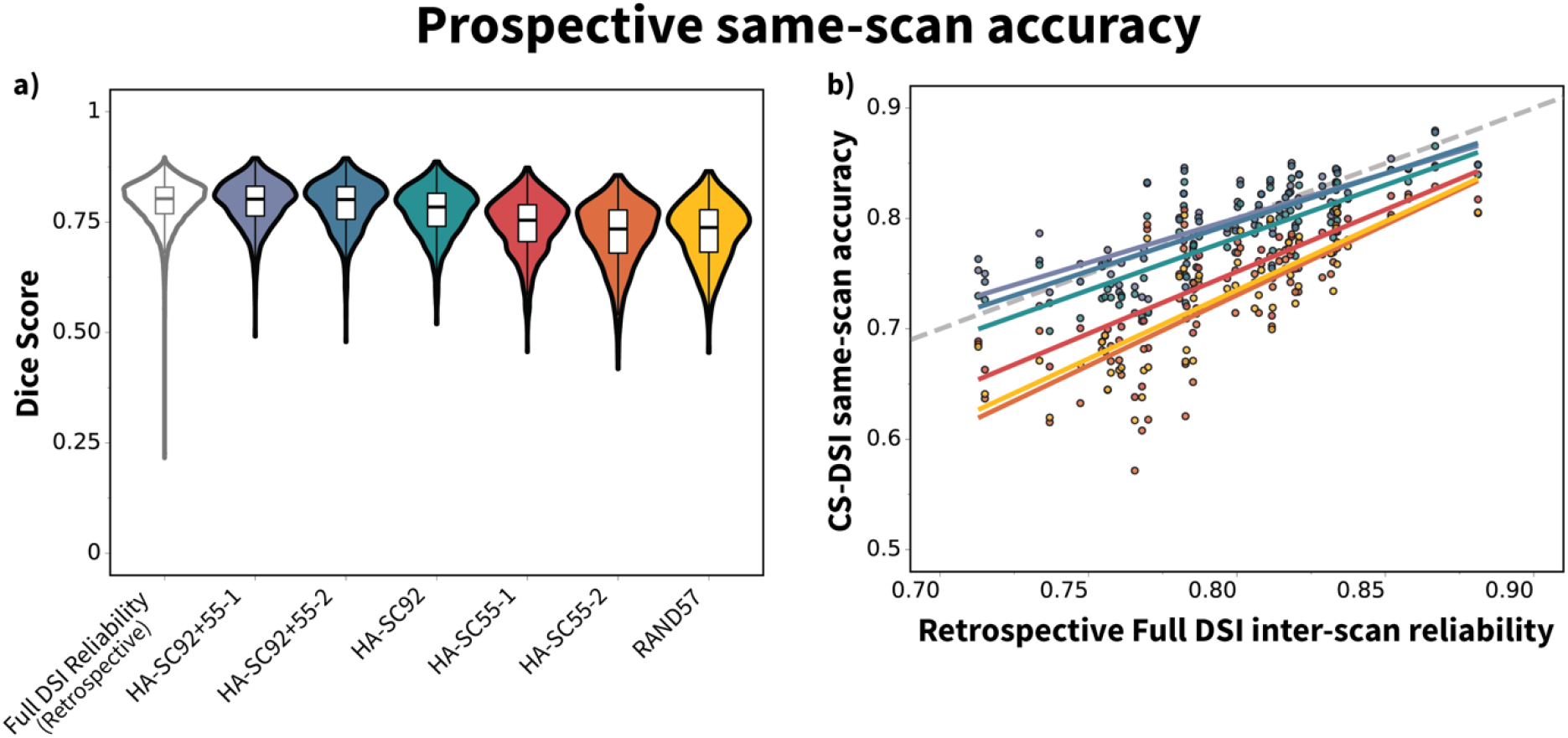
CS-DSI accuracy of bundle segmentation can be replicated in a dataset where CS-DSI images are prospectively acquired. a) Dice scores between most CS-segmented bundles and combined DSI-segmented bundles are similar to the inter-scan reliability of full DSI bundles from the retrospective data. Violins represent distributions of Dice scores across all participants and bundles for a given scheme. b) Median distribution of the accuracy for each CS-DSI scheme is highly correlated with the median distribution of the full DSI inter-scan reliability from the retrospective data. Here, each point on the scatter plot represents the median of Dice scores for a single bundle across participants. The gray dashed lined denotes x=y.

Finally, we evaluated the accuracy of voxel-wise NQA, GFA, and ISO maps in the prospective data. We found that the maps derived from CS-DSI images were all highly correlated with maps derived from the full DSI image. Here too, the distribution of Pearson correlation coefficients between the CS-DSI derived scalars and the full DSI derived scalars was comparable with inter-scan reliability of the full DSI scheme (from the retrospective data) for almost all scalars studied (**Figure 10**). This replication of our results in an independent dataset that was prospectively acquired on a different scanner further bolster confidence in our findings and underscore the utility of CS-DSI.

**Figure 10:**
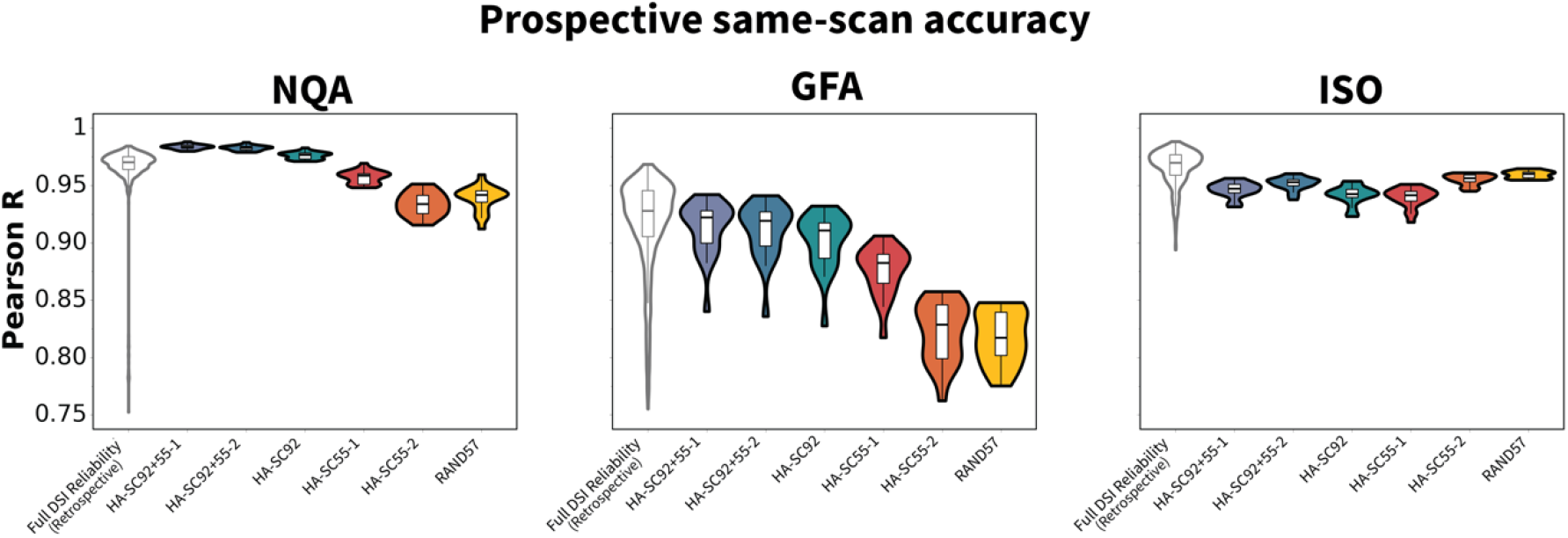
CS-DSI accuracy of scalar map generation can be replicated in a dataset where CS-DSI data is prospectively acquired. Pearson correlations of the voxel-wise scalars between the CS schemes and the combined scheme in the prospective dataset are similar to the full DSI inter-scan reliability from the retrospective data.

## 4 Discussion

CS-DSI has been shown to be a promising strategy to reduce DSI acquisition time in post-mortem and animal studies. However, it is unclear whether this approach is a viable alternative to full DSI in conducting rigorous studies of human brain white matter *in vivo*. In this study, we examined the capacity for six different CS-DSI schemes to non-invasively quantify macrostructural and microstructural white matter properties in the human brain. We consistently found that both CS-DSI derived bundle segmentations and voxel-wise scalars were comparable in accuracy and reliability to those derived from full DSI images. Our results demonstrate that CS-DSI is an efficient and robust diffusion imaging technique that can accurately and reliably quantify white matter in a fraction of the acquisition time required by a full DSI scan, underscoring its immense clinical and research potential.

There are many approaches to construct CS schemes for dMRI. In this paper, we primarily examined HA-SC schemes with different *q*-space sampling densities, and a random scheme (RAND57) designed to enable concatenation to the full DSI scheme. The HA-SC scheme ensured a homogenous angular covering of the *q*-space, and had been previously shown to provide the most efficient and robust reconstruction of the EAP in simulated data (Paquette et al., 2015). We observed that all HA-SC schemes performed better than the RAND57 scheme, possibly because the homogenous distribution of subsampled points allowed for more effective extrapolation of the unacquired points in *q*-space. We also consistently observed that HA-SC accuracy and reliability scaled with the number of directions sampled, with the longer, more densely sampled schemes performing better.

To further contextualize the performance of our validity measures, we compared them against the inter-scan reliability of the full DSI scheme. Inter-scan reliability reflects test-retest differences, which are inevitable in neuroimaging, even when using longer acquisition schemes. We reasoned that if differences between CS-DSI derivatives and full DSI derivatives were similar to the test-retest differences in full DSI derivatives, the corresponding CS-DSI scans could be considered as viable alternatives to full DSI. We showed that validity measures of select CS-DSI schemes were similar to full DSI reliability for both bundle segmentations and scalar quantification. Impressively, while the HA-SC92 scheme took only around a third of the acquisition time required for full DSI, this reduction in acquisition time had no negative impact on its data quality: HA-SC92 accuracies and reliabilities were not statistically different from full DSI reliabilities for almost all derivatives examined. Though HA-SC92+55-1 and HA-SC92+55-2 also performed very well, the HA-SC92 scheme had an acquisition time of only 7.4 minutes. These results suggest that the information lost from subsampling the *q*-space in the denser HA-SC schemes was minimal and comparable with mere test-retest differences of the full DSI. This key finding demonstrates that the HA-SC92 scheme can be successfully used as an alternative for full DSI with very few trade-offs for the derived measures we examined.

While the more sparsely sampled schemes with shorter scan times had validity metrics that were often significantly different from the full DSI reliability, we found that these differences were very small, especially for the HA-SC55-1 and HA-SC55-2 schemes. All bundles segmented by HA-SC55-1 and HA-SC55-2 were still qualitatively comparable to those segmented by full DSI with the correct shapes, and minimal spurious streamlines. Moreover, for bundles that were more reliably segmented by full DSI (like the Corpus Callosum or the Parietal Aslant), even the more sparsely sampled schemes had very high validity measures that were not statistically different from full DSI reliability. These results suggest that scientific endeavors focused on these specific bundles could safely adopt a 4-minute CS-DSI scan. This feature of the sparser CS-DSI schemes is particularly valuable in contexts where scan time needs to be as low as possible, like in clinical care or in research of populations where extended scans are impractical (e.g., children, clinical populations).

These differences in HA-SC validity and full DSI reliability were even smaller when computing the voxel-wise scalars. The median Pearson correlation between HA-SC and full DSI scalar maps was consistently above 0.85 for all scalars examined. This finding is particularly promising as it suggests that CS-DSI has the capacity to successfully detect specific and localized microstructural changes. The high correlation between HA-SC and full DSI scalars suggests that factors that are associated with full DSI scalars would also be associated with HA-SC scalars, allowing us to detect the same properties in a fraction of the scan time. Future studies correlating HA-SC derivatives with factors of interest like age, cognition, or clinical status, and comparing these relationships with those from full DSI will be able to further ascertain CS-DSI’s utility in these contexts.

Despite being slightly longer, the RAND57 scheme did not perform as well as any of the HA-SC schemes, and showed significant differences against full DSI reliability in almost all its validity measures, when generating both bundles and scalar maps. These findings indicate that not all CS-DSI schemes are equivalent, and that *q*-space subsampling must be performed strategically to be effective. This observation also highlights the significance of our HA-SC results: while not all CS-DSI schemes are effective, we identified a sampling strategy that can approximate full DSI performance in a fraction of the scan time.

Not only can HA-SC schemes be an effective alternative to full DSI, but they might also be able to improve all studies leveraging diffusion imaging. Single shell diffusion schemes continue to be the most popular DWI acquisitions, especially in clinically relevant endeavors, given their short scan times. The HA-SC55 schemes have similar acquisition times as a traditional single shell scheme required to generate tensors (Soares et al., 2013). However, these CS-DSI schemes can resolve crossing fibers and quantify more specific microstructural properties, unlike single shell schemes. A recent study even showed that CS-DSI was better than DTI at visualizing motor and language white matter tracks in patients with brain tumors (Young et al., 2017), further underscoring its clinical potential. Moreover, CS-DSI schemes can be effectively extrapolated into other HARDI schemes like multi-shell schemes, and perhaps derive even more scalars sensitive to microstructure (Tobisch et al., 2018), a feature not afforded by other fast schemes.

This study has several limitations that should be acknowledged. First, the retrospective CS-DSI images were subsampled from the full DSI dataset *after* preprocessing. Within each scan session, the motion correction on the subsampled volumes could also leverage from volumes that were not a part of the examined scheme, giving the retrospectively synthesized images an advantage not available to prospectively acquired CS-DSI data. Given this advantage, along with the absence of inter-scan noise, it is unsurprising that same-scan accuracy was higher than full DSI inter-scan reliability for some of the denser CS-DSI schemes. We addressed this problem by both calculating inter-scan accuracy, as well as replicating these results in prospectively acquired CS-DSI data. Moreover, while our results demonstrate minimal differences in validity measures between CS-DSI and full DSI derivatives, we cannot gauge the practical consequences of these differences. While beyond the scope of this paper, future studies examining the relationship between subsampling and correlations with metrics of interest (like age, cognition, or pathological outcomes) will help answer this question. Third, we used SHORE basis functions with L2-regularization to perform compressed sensing. While L2-regularization has been shown to be an effective way to extrapolate the full DSI signal from the CS-DSI data (Merlet, 2010; Merlet and Deriche, 2013), there are many other CS-DSI strategies that use other basis functions (discrete cosine transform, discrete wavelet transform (Almasri et al., 2020; Hong et al., 2011; Lai et al., 2016)) or approaches. Recent studies have demonstrated that adaptive dictionary-based approaches can substantially improve reconstruction (Bilgic et al., 2013, 2012; Jones et al., 2021). While some of these improved methods may require more computational resources, they have the potential to further increase the bundle and scalar derivation validity of even the sparser CS-DSI schemes. Investigating these algorithms using the framework presented here may further improve white matter measures derived from CS-DSI acquisitions.

In conclusion, we demonstrated that CS-DSI can provide an accurate and reliable characterization of white matter properties in human brains *in vivo*, in a fraction of the scan time required for full DSI. This reduction in scan time will allow CS-DSI to be added to more routine neuroimaging protocols in large-scale research studies.

## ACKNOWLEDGMENTS

This study was supported by grants from the National Institutes of Health: R01MH112847 (R.T.S., T.D.S.), R01MH120482 (T.D.S.), R37MH125829 (T.D.S. & D.A.F), R01EB022573 (T.D.S.), R01MH113550 (T.D.S.), R01MH123550 (R.T.S.), RF1MH116920 (T.D.S.), K99MH127293 (B.L.), R01MH119185 (D.R.R), R01MH120174 (D.R.R.) and the U.S. Army Research Office and accomplished under cooperative agreement W911NF-19-2-0026 and contract W911NF-09-D-0001 for the Institute for Collaborative Biotechnologies. V.J.S. was supported by a National Science Foundation Graduate Research Fellowship (DGE-1845298). Additional support was provided by the AE Foundation and the Penn/CHOP Lifespan Brain Institute.

## Declaration of Interest

The data in this manuscript were collected in compliance with ethical standards. These data have not been published previously and are not under consideration for publication elsewhere. All authors have contributed significantly to this manuscript and have approved this submission. All authors have no conflicts of interests in the conduct or reporting of this research.

## Notes

### Competing Interest Statement

The authors have declared no competing interest.

## References

Adluru, G., Dibella, E.V.R., 2008. Reordering for improved constrained reconstruction from undersampled k-space data. Int. J. Biomed. Imaging 2008, 341684. https://doi.org/10.1155/2008/341684

Almasri, N., Sadhu, A., Ray Chaudhuri, S., 2020. Toward Compressed Sensing of Structural Monitoring Data Using Discrete Cosine Transform. J. Comput. Civ. Eng. 34, 04019041. https://doi.org/10.1061/(ASCE)CP.1943-5487.0000855

Assaf, Y., Basser, P.J., 2005. Composite hindered and restricted model of diffusion (CHARMED) MR imaging of the human brain. NeuroImage 27, 48–58. https://doi.org/10.1016/j.neuroimage.2005.03.042

Baraniuk, R., Steeghs, P., 2007. Compressive Radar Imaging, in: 2007 IEEE Radar Conference. Presented at the 2007 IEEE Radar Conference, pp. 128–133. https://doi.org/10.1109/RADAR.2007.374203

Benjamini, Y., Hochberg, Y., 1995. Controlling the False Discovery Rate: A Practical and Powerful Approach to Multiple Testing. J. R. Stat. Soc. Ser. B Methodol. 57, 289–300.

Berger, C.R., Zhou, S., Preisig, J.C., Willett, P., 2010. Sparse Channel Estimation for Multicarrier Underwater Acoustic Communication: From Subspace Methods to Compressed Sensing. IEEE Trans. Signal Process. 58, 1708–1721. https://doi.org/10.1109/TSP.2009.2038424

Bilgic, B., Chatnuntawech, I., Setsompop, K., Cauley, S.F., Yendiki, A., Wald, L.L., Adalsteinsson, E., 2013. Fast dictionary-based reconstruction for diffusion spectrum imaging. IEEE Trans. Med. Imaging 32, 2022–2033. https://doi.org/10.1109/TMI.2013.2271707

Bilgic, B., Setsompop, K., Cohen-Adad, J., Wedeen, V., Wald, L.L., Adalsteinsson, E., 2012. Accelerated Diffusion Spectrum Imaging with Compressed Sensing using Adaptive Dictionaries. Med. Image Comput. Comput.-Assist. Interv. MICCAI Int. Conf. Med. Image Comput. Comput.-Assist. Interv. 15, 1–9.

Bobin, J., Starck, J.-L., 2009. Compressed sensing in astronomy and remote sensing: a data fusion perspective, in: Goyal, V.K., Papadakis, M., Van De Ville, D. (Eds.),. Presented at the SPIE Optical Engineering + Applications, San Diego, CA, p. 74460I. https://doi.org/10.1117/12.830633

Callaghan, P.T., Eccles, C.D., Xia, Y., 1988. NMR microscopy of dynamic displacements: k-space and q-space imaging. J. Phys. [E] 21, 820. https://doi.org/10.1088/0022-3735/21/8/017

Candes, E.J., Wakin, M.B., 2008. An Introduction To Compressive Sampling. IEEE Signal Process. Mag. 25, 21–30. https://doi.org/10.1109/MSP.2007.914731

Carroll, M.K., Cecchi, G.A., Rish, I., Garg, R., Rao, A.R., 2009. Prediction and interpretation of distributed neural activity with sparse models. NeuroImage 44, 112–122. https://doi.org/10.1016/j.neuroimage.2008.08.020

Cheng, J., Merlet, S., Caruyer, E., Ghosh, A., Jiang, T., Deriche, R., 2011. Compressive Sensing Ensemble Average Propagator Estimation via L1 Spherical Polar Fourier Imaging. Presented at the MICCAI Workshop on Computational Diffusion MRI - CDMRI’11.

Cieslak, M., Cook, P.A., Tapera, T.M., Radhakrishnan, H., Elliott, M., Roalf, D.R., Oathes, D.J., Bassett, D.S., Tisdall, M.D., Rokem, A., Grafton, S.T., Satterthwaite, T.D., 2022. Diffusion MRI Head Motion Correction Methods are Highly Accurate but Impacted by Denoising and Sampling Scheme. https://doi.org/10.1101/2022.07.21.500865

Cieslak, M., Cook, P.A., He, X. et al., 2021. QSIPrep: an integrative platform for preprocessing and reconstructing diffusion MRI data. Nature Methods 18, 775–778 https://doi.org/10.1038/s41592-021-01185-5

Cieslak, M., Meiring, W., Brennan, T., Greene, C., Volz, L.J., Vettel, J.M., Suri, S., Grafton, S.T., 2018. Compositional measures of diffusion anisotropy and asymmetry, in: 2018 IEEE 15th International Symposium on Biomedical Imaging (ISBI 2018). Presented at the 2018 IEEE 15th International Symposium on Biomedical Imaging (ISBI 2018), pp. 123–126. https://doi.org/10.1109/ISBI.2018.8363537

Donoho, D.L., 2006. Compressed sensing. IEEE Trans. Inf. Theory 52, 1289–1306. https://doi.org/10.1109/TIT.2006.871582

Gorgolewski, K., Burns, C., Madison, C., Clark, D., Halchenko, Y., Waskom, M., Ghosh, S., 2011. Nipype: A Flexible, Lightweight and Extensible Neuroimaging Data Processing Framework in Python. Front. Neuroinformatics 5.

Gorgolewski, K.J., Esteban, O., Ellis, D.G., Notter, M.P., Ziegler, E., Johnson, H., Hamalainen, C., Yvernault, B., Burns, C., Manhães-Savio, A., Jarecka, D., Markiewicz, C.J., Salo, T., Clark, Daniel, Waskom, M., Wong, J., Modat, M., Dewey, B.E., Clark, M.G., Dayan, M., Loney, F., Madison, C., Gramfort, A., Keshavan, A., Berleant, S., Pinsard, B., Goncalves, M., Clark, Dav, Cipollini, B., Varoquaux, G., Wassermann, D., Rokem, A., Halchenko, Y.O., Forbes, J., Moloney, B., Malone, I.B., Hanke, M., Mordom, D., Buchanan, C., Pauli, W.M., Huntenburg, J.M., Horea, C., Schwartz, Y., Tungaraza, R., Iqbal, S., Kleesiek, J., Sikka, S., Frohlich, C., Kent, J., Perez-Guevara, M., Watanabe, A., Welch, D., Cumba, C., Ginsburg, D., Eshaghi, A., Kastman, E., Bougacha, S., Blair, R., Acland, B., Gillman, A., Schaefer, A., Nichols, B.N., Giavasis, S., Erickson, D., Correa, C., Ghayoor, A., Küttner, R., Haselgrove, C., Zhou, D., Craddock, R.C., Haehn, D., Lampe, L., Millman, J., Lai, J., Renfro, M., Liu, S., Stadler, J., Glatard, T., Kahn, A.E., Kong, X.-Z., Triplett, W., Park, A., McDermottroe, C., Hallquist, M., Poldrack, R., Perkins, L.N., Noel, M., Gerhard, S., Salvatore, J., Mertz, F., Broderick, W., Inati, S., Hinds, O., Brett, M., Durnez, J., Tambini, A., Rothmei, S., Andberg, S.K., Cooper, G., Marina, A., Mattfeld, A., Urchs, S., Sharp, P., Matsubara, K., Geisler, D., Cheung, B., Floren, A., Nickson, T., Pannetier, N., Weinstein, A., Dubois, M., Arias, J., Tarbert, C., Schlamp, K., Jordan, K., Liem, F., Saase, V., Harms, R., Khanuja, R., Podranski, K., Flandin, G., Papadopoulos Orfanos, D., Schwabacher, I., McNamee, D., Falkiewicz, M., Pellman, J., Linkersdörfer, J., Varada, J., Pérez-García, F., Davison, A., Shachnev, D., Ghosh, S., 2017. Nipype: a flexible, lightweight and extensible neuroimaging data processing framework in Python. 0.13.1. https://doi.org/10.5281/zenodo.581704

Gramfort, A., Poupon, C., Descoteaux, M., 2012. Sparse DSI: Learning DSI Structure for Denoising and Fast Imaging, in: Ayache, N., Delingette, H., Golland, P., Mori, K. (Eds.), Medical Image Computing and Computer-Assisted Intervention – MICCAI 2012, Lecture Notes in Computer Science. Springer, Berlin, Heidelberg, pp. 288–296. https://doi.org/10.1007/978-3-642-33418-4_36

He, L., Chang, T.-C., Osher, S., Fang, T., Speier, P., 2007. MR image reconstruction from undersampled data by using the iterative refinement procedure. PAMM 7, 1011207– 1011208. https://doi.org/10.1002/pamm.200700776

Herman, M.A., Strohmer, T., 2009. High-Resolution Radar via Compressed Sensing. IEEE Trans. Signal Process. 57, 2275–2284. https://doi.org/10.1109/TSP.2009.2014277

Hong, M., Yu, Y., Wang, H., Liu, F., Crozier, S., 2011. Compressed sensing MRI with singular value decomposition-based sparsity basis. Phys. Med. Biol. 56, 6311. https://doi.org/10.1088/0031-9155/56/19/010

Jaspan, O.N., Fleysher, R., Lipton, M.L., 2015. Compressed sensing MRI: a review of the clinical literature. Br. J. Radiol. 88, 20150487. https://doi.org/10.1259/bjr.20150487

Jones, D.K., Horsfield, M.A., Simmons, A., 1999. Optimal strategies for measuring diffusion in anisotropic systems by magnetic resonance imaging. Magn. Reson. Med. 42, 515–525.

Jones, R., Maffei, C., Augustinack, J., Fischl, B., Wang, H., Bilgic, B., Yendiki, A., 2021. High-fidelity approximation of grid- and shell-based sampling schemes from undersampled DSI using compressed sensing: Post mortem validation. NeuroImage 244, 118621. https://doi.org/10.1016/j.neuroimage.2021.118621

Lai, Z., Qu, X., Liu, Y., Guo, D., Ye, J., Zhan, Z., Chen, Z., 2016. Image reconstruction of compressed sensing MRI using graph-based redundant wavelet transform. Med. Image Anal., Discrete Graphical Models in Biomedical Image Analysis 27, 93–104. https://doi.org/10.1016/j.media.2015.05.012

Lustig, M., Donoho, D., Pauly, J.M., 2007. Sparse MRI: The application of compressed sensing for rapid MR imaging. Magn. Reson. Med. 58, 1182–1195. https://doi.org/10.1002/mrm.21391

Lustig, M., Donoho, D.L., Santos, J.M., Pauly, J.M., 2008. Compressed Sensing MRI. IEEE Signal Process. Mag. 25, 72–82. https://doi.org/10.1109/MSP.2007.914728

Menzel, M.I., Tan, E.T., Khare, K., Sperl, J.I., King, K.F., Tao, X., Hardy, C.J., Marinelli, L., 2011a. Accelerated diffusion spectrum imaging in the human brain using compressed sensing. Magn. Reson. Med. 66, 1226–1233. https://doi.org/10.1002/mrm.23064

Menzel, M.I., Tan, E.T., Khare, K., Sperl, J.I., King, K.F., Tao, X., Hardy, C.J., Marinelli, L., 2011b. Accelerated diffusion spectrum imaging in the human brain using compressed sensing. Magn. Reson. Med. 66, 1226–1233. https://doi.org/10.1002/mrm.23064

Merlet, S., Deriche, R., 2010 Compressed Sensing for Accelerated EAP Recovery in Diffusion MRI. MICCAI, Pekin, China. pp.Page 14. ffinria-00536278f

Merlet, S., 2013. Compressive Sensing in diffusion MRI. Medical Imaging. Université Nice Sophia Antipolis, English. ffNNT : ff. fftel-00908369v1f

Merlet, S.L., Deriche, R., 2013. Continuous diffusion signal, EAP and ODF estimation via Compressive Sensing in diffusion MRI. Med. Image Anal. 17, 556–572. https://doi.org/10.1016/j.media.2013.02.010

Michailovich, O., Rathi, Y., Dolui, S., 2011. Spatially regularized compressed sensing for high angular resolution diffusion imaging. IEEE Trans. Med. Imaging 30, 1100–1115. https://doi.org/10.1109/TMI.2011.2142189

Naeyaert, M., Aelterman, J., Van Audekerke, J., Golkov, V., Cremers, D., Pižurica, A., Sijbers, J., Verhoye, M., 2021. Accelerating in vivo fast spin echo high angular resolution diffusion imaging with an isotropic resolution in mice through compressed sensing. Magn. Reson. Med. 85, 1397–1413. https://doi.org/10.1002/mrm.28520

Nakuci, J., Wasylyshyn, N., Cieslak, M., Elliot, J.C., Bansal, K., Giesbrecht, B., Grafton, S.T., Vettel, J.M., Garcia, J.O., Muldoon, S.F., 2022. Within- and between-subject reproducibility and variability in multi-modal, longitudinal brain networks (preprint). Neuroscience. https://doi.org/10.1101/2022.05.03.490544

Ozarslan, E., Koay, C., Shepherd, T.M., Blackb, S.J., Basser, P.J., 2013. Simple harmonic oscillator based reconstruction and estimation for three-dimensional q-space MRI.

Paquette, M., Merlet, S., Gilbert, G., Deriche, R., Descoteaux, M., 2015. Comparison of sampling strategies and sparsifying transforms to improve compressed sensing diffusion spectrum imaging. Magn. Reson. Med. 73, 401–416. https://doi.org/10.1002/mrm.25093

Pu, L., Trouard, T.P., Ryan, L., Huang, C., Altbach, M.I., Bilgin, A., 2011. Model-based compressive diffusion tensor imaging, in: 2011 IEEE International Symposium on Biomedical Imaging: From Nano to Macro. Presented at the 2011 IEEE International Symposium on Biomedical Imaging: From Nano to Macro, pp. 254–257. https://doi.org/10.1109/ISBI.2011.5872400

Ramirez-Manzanares, A., Rivera, M., Vemuri, B.C., Carney, P., Mareci, T., 2007. Diffusion basis functions decomposition for estimating white matter intravoxel fiber geometry. IEEE Trans. Med. Imaging 26, 1091–1102. https://doi.org/10.1109/TMI.2007.900461

Schilling, K.G., Yeh, F.-C., Nath, V., Hansen, C., Williams, O., Resnick, S., Anderson, A.W., Landman, B.A., 2019. A fiber coherence index for quality control of B-table orientation in diffusion MRI scans. Magn. Reson. Imaging 58, 82–89. https://doi.org/10.1016/j.mri.2019.01.018

Soares, J., Marques, P., Alves, V., Sousa, N., 2013. A hitchhiker’s guide to diffusion tensor imaging. Front. Neurosci. 7.

Tobisch, A., Stirnberg, R., Harms, R.L., Schultz, T., Roebroeck, A., Breteler, M.M.B., Stöcker, T., 2018. Compressed Sensing Diffusion Spectrum Imaging for Accelerated Diffusion Microstructure MRI in Long-Term Population Imaging. Front. Neurosci. 12.

Tristán-Vega, A., Westin, C.-F., 2011. Probabilistic ODF Estimation from Reduced HARDI Data with Sparse Regularization, in: Fichtinger, G., Martel, A., Peters, T. (Eds.), Medical Image Computing and Computer-Assisted Intervention – MICCAI 2011, Lecture Notes in Computer Science. Springer, Berlin, Heidelberg, pp. 182–190. https://doi.org/10.1007/978-3-642-23629-7_23

Tuch, D.S., Reese, T.G., Wiegell, M.R., Makris, N., Belliveau, J.W., Wedeen, V.J., 2002. High angular resolution diffusion imaging reveals intravoxel white matter fiber heterogeneity. Magn. Reson. Med. 48, 577–582. https://doi.org/10.1002/mrm.10268

Wang, C., Holly, L.T., Oughourlian, T., Yao, J., Raymond, C., Salamon, N., Ellingson, B.M., 2021. Detection of cerebral reorganization associated with degenerative cervical myelopathy using diffusion spectral imaging (DSI). J. Clin. Neurosci. 86, 164–173. https://doi.org/10.1016/j.jocn.2021.01.011

Wang, N., White, L.E., Qi, Y., Cofer, G., Johnson, G.A., 2020. Cytoarchitecture of the mouse brain by high resolution diffusion magnetic resonance imaging. NeuroImage 216, 116876. https://doi.org/10.1016/j.neuroimage.2020.116876

Wedeen, V.J., Hagmann, P., Tseng, W.Y.I., Reese, T.G., Weisskoff, R.M., 2005. Mapping complex tissue architecture with DSI magnetic resonance imaging. Magn. Reson. Med. 54, 1377–1385.

Wedeen, Van J., Hagmann, P., Tseng, W.-Y.I., Reese, T.G., Weisskoff, R.M., 2005. Mapping complex tissue architecture with diffusion spectrum magnetic resonance imaging. Magn. Reson. Med. 54, 1377–1386. https://doi.org/10.1002/mrm.20642

Wedeen, V.J., Wang, R.P., Schmahmann, J.D., Benner, T., Tseng, W.Y.I., Dai, G., Pandya, D.N., Hagmann, P., D’Arceuil, H., de Crespigny, A.J., 2008. Diffusion spectrum magnetic resonance imaging (DSI) tractography of crossing fibers. NeuroImage 41, 1267–1277. https://doi.org/10.1016/j.neuroimage.2008.03.036

Wiaux, Y., Jacques, L., Puy, G., Scaife, A.M.M., Vandergheynst, P., 2009. Compressed sensing imaging techniques for radio interferometry. Mon. Not. R. Astron. Soc. 395, 1733– 1742. https://doi.org/10.1111/j.1365-2966.2009.14665.x

Ye, W., Vemuri, B.C., Entezari, A., 2012. An over-complete dictionary based regularized reconstruction of a field of ensemble average propagators, in: 2012 9th IEEE International Symposium on Biomedical Imaging (ISBI). Presented at the 2012 9th IEEE International Symposium on Biomedical Imaging (ISBI), pp. 940–943. https://doi.org/10.1109/ISBI.2012.6235711

Yeh, F.-C., 2020. Shape analysis of the human association pathways. NeuroImage 223, 117329. https://doi.org/10.1016/j.neuroimage.2020.117329

Yeh, F.-C., Panesar, S., Barrios, J., Fernandes, D., Abhinav, K., Meola, A., Fernandez-Miranda, J.C., 2019. Automatic Removal of False Connections in Diffusion MRI Tractography Using Topology-Informed Pruning (TIP). Neurother. J. Am. Soc. Exp. Neurother. 16, 52–58. https://doi.org/10.1007/s13311-018-0663-y

Yeh, F.-C., Panesar, S., Fernandes, D., Meola, A., Yoshino, M., Fernandez-Miranda, J.C., Vettel, J.M., Verstynen, T., 2018. Population-averaged atlas of the macroscale human structural connectome and its network topology. NeuroImage 178, 57–68. https://doi.org/10.1016/j.neuroimage.2018.05.027

Yeh, F.-C., Verstynen, T.D., Wang, Y., Fernández-Miranda, J.C., Tseng, W.-Y.I., 2013. Deterministic Diffusion Fiber Tracking Improved by Quantitative Anisotropy. PLOS ONE 8, e80713. https://doi.org/10.1371/journal.pone.0080713

Yeh, F.-C., Vettel, J.M., Singh, A., Poczos, B., Grafton, S.T., Erickson, K.I., Tseng, W.-Y.I., Verstynen, T.D., 2016. Quantifying Differences and Similarities in Whole-Brain White Matter Architecture Using Local Connectome Fingerprints. PLOS Comput. Biol. 12, e1005203. https://doi.org/10.1371/journal.pcbi.1005203

Yeh, F.-C., Wedeen, V.J., Tseng, W.-Y.I., 2010. Generalized $ q$-Sampling Imaging. IEEE Trans. Med. Imaging 29, 1626–1635. https://doi.org/10.1109/TMI.2010.2045126

Yoshiura, T., Wu, O., Zaheer, A., Reese, T.G., Sorensen, A.G., 2001. Highly diffusion-sensitized MRI of brain: dissociation of gray and white matter. Magn. Reson. Med. 45, 734– 740. https://doi.org/10.1002/mrm.1100

Young, R.J., Tan, E.T., Peck, K.K., Jenabi, M., Karimi, S., Brennan, N., Rubel, J., Lyo, J., Shi, W., Zhang, Z., Prastawa, M., Liu, X., Sperl, J.I., Fatovic, R., Marinelli, L., Holodny, A.I., 2017. Comparison of compressed sensing diffusion spectrum imaging and diffusion tensor imaging in patients with intracranial masses. Magn. Reson. Imaging 36, 24–31. https://doi.org/10.1016/j.mri.2016.10.001

Zhang, H., Wang, Yong, Lu, T., Qiu, B., Tang, Y., Ou, S., Tie, X., Sun, C., Xu, K., Wang, Yibao, 2013. Differences between generalized q-sampling imaging and diffusion tensor imaging in the preoperative visualization of the nerve fiber tracts within peritumoral edema in brain. Neurosurgery 73, 1044–1053; discussion 1053. https://doi.org/10.1227/NEU.0000000000000146

