## Supplementary for "Establishing the Validity of Compressed Sensing Diffusion Spectrum Imaging"

Supplementary Material

**Permutation testing for the retrospective data**

To assess CS-DSI validity, we compared the accuracy and reliability distributions of these schemes to the reliability distribution of the full DSI data. As each of these distributions had a very large number of data points and were not independent of each other, traditional parametric statistics were not appropriate to quantify statistical differences. Accordingly, to determine if the distributions of CS-DSI validity metrics were significantly different from full DSI reliability, we utilized permutation testing. Permutation testing could only be performed on the retrospective dataset, as full DSI reliability could not be calculated on the prospective data due to the absence of multiple scan sessions. Accuracy and reliability required different designs and are described below. For each validity metric, permutation tests were designed identically for bundles and scalars. Each CS-DSI scheme was tested independently against the full DSI scheme, by comparing distributions of CS-DSI validity metrics with distributions of full DSI reliability. For each CS-DSI scheme, FDR-corrected two-sided *p*-values were calculated per bundle and per scalar.

**CS-DSI accuracy vs full DSI reliability statistics**

For each participant, there were 8 values (either Dice scores or Pearson Rs) for same-scan accuracy (1 for each of the 8 sessions), 56 values for inter-scan accuracy ($8P2$ for each pairwise combination of sessions), and 28 values for inter-scan reliability ($8C2$ for each pairwise combination of sessions). Each of the CS-DSI accuracy distributions were compared against the full DSI reliability distribution for each bundle and diffusion scheme. Permutation testing for both same-scan and inter-scan accuracy was performed the same way. The difference between distributions was calculated by subtracting the median of the CS-DSI accuracy distribution from the median of the full DSI reliability distribution for each participant. We then calculated the median of these differences across participants, to yield an overall median for each bundle or scalar metric. This was referred to as the “observed” median difference for that CS-DSI scheme’s bundle or scalar.

To determine if the observed median difference was statistically significant, we generated a null distribution for each CS-DSI scheme’s measure. We randomly permuted the labels (CS-DSI accuracy or full DSI reliability) of these values while maintaining the proportions of these labels (28:8 for same-scan accuracy and 28:56 for inter-scan accuracy) for each participant. We then calculated the median difference as above, and repeated this procedure 1000 times per participant. This set of permuted median differences formed our null distribution. To get an overall null distribution per bundle or scalar metric, we concatenated the null distributions across participants. Two-sided *p*-values were then calculated based on the position of the observed median within the overall null distribution. A single *p*-value was calculated per bundle or voxel-wise scalar. These *p*-values were then corrected for multiple comparisons (against 56 comparisons for the bundles and 3 comparisons for the voxel-wise scalars) using the Benjamini/Hochberg False Discovery Rate method (Benjamini and Hochberg, 1995) (**Supplementary Tables 2, 4, 6, and 7)**.

**CS-DSI reliability vs full DSI reliability statistics**

For this test, as the same validity metric was compared between the CS-DSI and full DSI schemes, we used a paired permutation testing design. For each pair of sessions within a participant, the difference between full DSI and CS-DSI reliability was calculated (Full DSI – CS-DSI). We then calculated the median of these differences across participants and session pairs to get the “observed” median difference between these distributions. For reliability, we generated our null distribution without breaking the integrity of the pair-wise relationships. For each pair of sessions, we randomly shuffled whether the difference in reliability would be multiplied by -1 or 1 i.e., whether the labels “Full DSI reliability” and “CS-DSI reliability” would be shuffled within that pair. We then calculated the median difference the same way as before, and repeated this shuffling 1000 times per participant. This set of shuffled median differences formed our null distribution. To arrive at an overall null distribution per bundle or scalar metric, we concatenated the null distributions across participants. Two-sided *p*-values were then calculated based on the position of the observed median within the overall null distribution. A single *p*-value was calculated per bundle or voxel-wise scalar. These *p*-values were then corrected for multiple comparisons (against 56 comparisons for the bundles and 3 comparisons for the voxel-wise scalars) using the Benjamini/Hochberg False Discovery Rate method (Benjamini and Hochberg, 1995) **(Supplementary Tables 5 and 8)**.

**Supplementary Tables**

| Bundle class | Corresponding Bundles |
| --- | --- |
| Aslant | Frontal Aslant (Left and Right),  Parietal Aslant (Left and Right) |
| Cingulum | Frontal Parahippocampal Cingulum (Left and Right),  Frontal Parietal Cingulum (Left and Right),  Parahippocampal Parietal Cingulum (Left and Right),  Parahippocampal Cingulum (Left* and Right*),  Paraolfactory Cingulum (Left and Right), |
| Corpus | Corpus Callosum Forceps Major,  Corpus Callosum Body,  Corpus Callosum Tapetum,  Corpus Callosum Forceps Minor |
| Corticos | Corticospinal Tract (Left and Right),  Anterior Corticostriatal Tract (Left and Right),  Posterior Corticostriatal Tract (Left and Right),  Superior Corticostriatal Tract (Left and Right) |
| Fasciculus | Arcuate Fasciculus (Left and Right),  Inferior Fronto-Occipital Fasciculus (Left and Right),  Inferior Longitudinal Fasciculus (Left and Right),  Middle Longitudinal Fasciculus (Left and Right),  Superior Longitudinal Fasciculus – 1 (Left and Right),  Superior Longitudinal Fasciculus – 2 (Left and Right),  Superior Longitudinal Fasciculus – 3 (Left and Right),  Uncinate Fasciculus (Left and Right),  Vertical Occipital Fasciculus (Left* and Right*) |
| Fornix | Fornix (Left* and Right) |
| Optic | Optic Radiation (Left and Right) |
| Reticular | Reticular Tract (Left and Right) |
| Thalamic | Anterior Thalamic Radiation (Left and Right),  Posterior Thalamic Radiation (Left and Right),  Superior Thalamic Radiation (Left and Right) |

**Supplementary Table 1:** List of white matter bundles segmented and examined. * indicates that segmentation failed for at least one attempt.

| CS-DSI scheme | Median difference between CS-DSI same-scan accuracy and full DSI reliability across all bundles | Number of bundles (out of 56) that show significant differences (FDR corrected p<0.05) between same-scan accuracy and full DSI reliability |
| --- | --- | --- |
| HA-SC92+55-1 | -0.013 | 0/56 |
| HA-SC92+55-2 | -0.017 | 0/56 |
| HA-SC92 | 0.002 | 0/56 |
| HA-SC55-1 | 0.039 | 52/56 |
| HA-SC55-2 | 0.058 | 53/56 |
| RAND57 | 0.074 | 56/56 |

**Supplementary Table 2:** Median differences in Dice scores and proportion of bundles that have significantly different (FDR corrected p<0,05) *same-scan accuracies* from full DSI inter-scan reliability for all CS-DSI schemes in the retrospective dataset.

| Bundle | Full DSI Reliability |
| --- | --- |
| Corpus Callosum Forceps Minor | 0.881 |
| Corpus Callosum Body | 0.867 |
| Parietal Aslant Tract (Right) | 0.858 |
| Corpus Callosum Forceps Major | 0.852 |
| Vertical Occipital Fasciculus (Right) | 0.837 |
| Corticostriatal Tract Superior (Right) | 0.834 |
| Corticostriatal Tract Anterior (Left) | 0.834 |
| Parietal Aslant Tract (Left) | 0.833 |
| Corticostriatal Tract Superior (Left) | 0.833 |
| Inferior Longitudinal Fasciculus (Left) | 0.832 |
| Superior Longitudinal Fasciculus3 (Left) | 0.832 |
| Frontal Aslant Tract (Left) | 0.829 |
| Corticospinal Tract (Right) | 0.821 |
| Cingulum Frontal Parietal (Right) | 0.821 |
| Corticostriatal Tract Posterior (Left) | 0.820 |
| Corticospinal Tract (Left) | 0.818 |
| Cingulum Frontal Parietal (Left) | 0.818 |
| Superior Longitudinal Fasciculus3 (Right) | 0.817 |
| Uncinate Fasciculus (Left) | 0.816 |
| Corticostriatal Tract Anterior (Right) | 0.814 |
| Corticostriatal Tract Posterior (Right) | 0.812 |
| Thalamic Radiation Anterior (Right) | 0.812 |
| Frontal Aslant Tract (Right) | 0.811 |
| Uncinate Fasciculus (Right) | 0.808 |
| Superior Longitudinal Fasciculus1 (Right) | 0.807 |
| Inferior Fronto Occipital Fasciculus (Left) | 0.806 |
| Superior Longitudinal Fasciculus2 (Right) | 0.801 |
| Inferior Longitudinal Fasciculus (Right) | 0.800 |
| Arcuate Fasciculus (Left) | 0.799 |
| Thalamic Radiation Superior (Left) | 0.797 |
| Thalamic Radiation Posterior (Left) | 0.787 |
| Inferior Fronto Occipital Fasciculus (Right) | 0.787 |
| Optic Radiation (Left) | 0.786 |
| Thalamic Radiation Posterior (Right) | 0.785 |
| Fornix (Right) | 0.785 |
| Thalamic Radiation Anterior (Left) | 0.783 |
| Cingulum Parahippocampal (Right) | 0.783 |
| Superior Longitudinal Fasciculus2 (Left) | 0.782 |
| Superior Longitudinal Fasciculus1 (Left) | 0.782 |
| Thalamic Radiation Superior (Right) | 0.781 |
| Middle Longitudinal Fasciculus (Left) | 0.770 |
| Vertical Occipital Fasciculus (Left) | 0.770 |
| Corpus Callosum Tapetum | 0.769 |
| Optic Radiation (Right) | 0.768 |
| Fornix (Left) | 0.766 |
| Reticular Tract (Left) | 0.761 |
| Middle Longitudinal Fasciculus (Right) | 0.760 |
| Reticular Tract (Right) | 0.758 |
| Cingulum Parahippocampal (Left) | 0.756 |
| Cingulum Parahippocampal Parietal (Right) | 0.756 |
| Cingulum Parahippocampal Parietal (Left) | 0.755 |
| Cingulum Frontal Parahippocampal (Right) | 0.747 |
| Cingulum Parolfactory (Right) | 0.737 |
| Cingulum Frontal Parahippocampal (Left) | 0.734 |
| Cingulum Parolfactory (Left) | 0.715 |
| Arcuate Fasciculus (Right) | 0.713 |

**Supplementary Table 3**:

Median inter-scan reliability of all bundles segmented by the full DSI scheme (from the retrospective dataset), in ascending order of reliability.

| CS-DSI scheme | Median difference between CS-DSI inter-scan accuracy and full DSI reliability across all bundles | Number of bundles (out of 56) that show significant differences (FDR corrected p<0.05) between inter-scan accuracy and full DSI reliability |
| --- | --- | --- |
| HA-SC92+55-1 | 0.024 | 56/56 |
| HA-SC92+55-2 | 0.022 | 53/56 |
| HA-SC92 | 0.034 | 55/56 |
| HA-SC55-1 | 0.060 | 56/56 |
| HA-SC55-2 | 0.074 | 56/56 |
| RAND57 | 0.089 | 56/56 |

**Supplementary Table 4:** Median differences in Dice scores and proportion of bundles that have significantly different (FDR corrected p<0.05) *inter-scan accuracies* from full DSI inter-scan reliability for all CS-DSI schemes in the retrospective dataset.

| CS-DSI scheme | Median difference between CS-DSI inter-scan reliability and full DSI reliability across all bundles | Number of bundles (out of 56) that show significant differences (FDR corrected p<0.05) between same-scan accuracy and full DSI reliability |
| --- | --- | --- |
| HA-SC92+55-1 | 0.008 | 0/56 |
| HA-SC92+55-2 | 0.009 | 0/56 |
| HA-SC92 | 0.018 | 0/56 |
| HA-SC55-1 | 0.038 | 0/56 |
| HA-SC55-2 | 0.039 | 0/56 |
| RAND57 | 0.063 | 0/56 |

**Supplementary Table 5:** Median differences in Dice scores and proportion of bundles that have significantly different (FDR corrected p<0.05) *inter-scan reliabilities* from full DSI for all CS-DSI schemes in the retrospective dataset.

**NQA**

| CS-DSI scheme | Median difference between CS-DSI same-scan accuracy and full DSI reliability | FDR corrected *p*-value |
| --- | --- | --- |
| **HA-SC92+55-1** | **-0.018** | **0.014** |
| **HA-SC92+55-2** | **-0.018** | **0.014** |
| **HA-SC92** | **-0.015** | **0.018** |
| HA-SC55-1 | 0.003 | 0.224 |
| **HA-SC55-2** | **0.022** | **0.015** |
| **RAND57** | **0.048** | **0.017** |

**GFA**

| CS-DSI scheme | Median difference between CS-DSI same-scan accuracy and full DSI reliability | FDR corrected *p*-value |
| --- | --- | --- |
| HA-SC92+55-1 | 0.009 | 0.374 |
| HA-SC92+55-2 | 0.008 | 0.398 |
| HA-SC92 | 0.014 | 0.237 |
| HA-SC55-1 | 0.046 | 0.172 |
| HA-SC55-2 | 0.065 | 0.057 |
| **RAND57** | **0.144** | **0.017** |

**ISO**

| CS-DSI scheme | Median difference between CS-DSI same-scan accuracy and full DSI reliability | FDR corrected *p*-value |
| --- | --- | --- |
| HA-SC92+55-1 | 0.010 | 0.207 |
| HA-SC92+55-2 | 0.012 | 0.154 |
| HA-SC92 | 0.011 | 0.167 |
| HA-SC55-1 | 0.010 | 0.224 |
| HA-SC55-2 | 0.015 | 0.060 |
| RAND57 | 0.005 | 0.323 |

**Supplementary Table 6:** Median differences in Pearson correlations and FDR corrected *p*-values for *same-scan accuracies* for all CS-DSI schemes and scalar metrics in the retrospective dataset. Rows in bold indicate statistically significant differences after correcting for multiple comparisons using the false discovery rate.

**NQA**

| CS-DSI scheme | Median difference between CS-DSI inter-scan accuracy and full DSI reliability | FDR corrected *p*-value |
| --- | --- | --- |
| **HA-SC92+55-1** | **0.009** | **0.001** |
| **HA-SC92+55-2** | **0.008** | **0.002** |
| **HA-SC92** | **0.012** | **<0.001** |
| **HA-SC55-1** | **0.029** | **0.002** |
| **HA-SC55-2** | **0.047** | **0.002** |
| **RAND57** | **0.075** | **0.001** |

**GFA**

| CS-DSI scheme | Median difference between CS-DSI inter-scan accuracy and full DSI reliability | FDR corrected *p*-value |
| --- | --- | --- |
| **HA-SC92+55-1** | **0.034** | **0.033** |
| **HA-SC92+55-2** | **0.034** | **0.033** |
| **HA-SC92** | **0.039** | **0.023** |
| **HA-SC55-1** | **0.068** | **0.006** |
| **HA-SC55-2** | **0.090** | **0.003** |
| **RAND57** | **0.170** | **0.001** |

**ISO**

| CS-DSI scheme | Median difference between CS-DSI inter-scan accuracy and full DSI reliability | FDR corrected *p*-value |
| --- | --- | --- |
| **HA-SC92+55-1** | **0.035** | **<0.001** |
| **HA-SC92+55-2** | **0.036** | **<0.001** |
| **HA-SC92** | **0.036** | **<0.001** |
| **HA-SC55-1** | **0.034** | **<0.001** |
| **HA-SC55-2** | **0.039** | **<0.001** |
| **RAND57** | **0.031** | **<0.001** |

**Supplementary Table 7:** Median differences in Pearson correlations and FDR corrected *p*-values for *inter-scan accuracies* for all CS-DSI schemes and scalar metrics in the retrospective dataset. Rows in bold indicate statistically significant differences after correcting for multiple comparisons using the false discovery rate.

**NQA**

| CS-DSI scheme | Median difference between CS-DSI inter-scan reliability and full DSI reliability | FDR corrected *p*-value |
| --- | --- | --- |
| HA-SC92+55-1 | 0.003 | 0.260 |
| HA-SC92+55-2 | 0.003 | 0.257 |
| HA-SC92 | 0.007 | 0.281 |
| HA-SC55-1 | 0.021 | 0.264 |
| HA-SC55-2 | 0.028 | 0.261 |
| RAND57 | 0.049 | 0.259 |

**GFA**

| CS-DSI scheme | Median difference between CS-DSI inter-scan reliability and full DSI reliability | FDR corrected *p*-value |
| --- | --- | --- |
| HA-SC92+55-1 | -0.042 | 0.214 |
| HA-SC92+55-2 | -0.042 | 0.204 |
| HA-SC92 | -0.038 | 0.200 |
| HA-SC55-1 | -0.019 | 0.216 |
| HA-SC55-2 | -0.009 | 0.261 |
| RAND57 | 0.026 | 0.259 |

**ISO**

| CS-DSI scheme | Median difference between CS-DSI inter-scan reliability and full DSI reliability | FDR corrected *p*-value |
| --- | --- | --- |
| HA-SC92+55-1 | 0.004 | 0.214 |
| HA-SC92+55-2 | 0.004 | 0.204 |
| HA-SC92 | 0.004 | 0.200 |
| HA-SC55-1 | 0.005 | 0.216 |
| HA-SC55-2 | 0.006 | 0.261 |
| RAND57 | 0.007 | 0.259 |

**Supplementary Table 8:** Median differences in Pearson correlations and FDR corrected *p*-values for *inter-scan reliabilities* for all CS-DSI schemes and scalar metrics in the retrospective dataset. Rows in bold indicate statistically significant differences after correcting for multiple comparisons using the false discovery rate.
